# PWK genetic background reprograms heterochromatin and malignant epithelial states in Neu-driven mammary tumors

**DOI:** 10.64898/2026.10.01.755936

**Authors:** Rui Geng, Ryann R. Ray, Jennifer B. Jacob, Benjamin L. Kidder

## Abstract

Genetic background can alter the latency and behavior of cancers initiated by the same oncogenic driver, but how inherited variation reshapes the regulatory landscape that permits tumor development remains poorly understood. The PWK genetic background accelerates mammary tumor onset in the NeuT model. We compared PWK NeuT and BALB NeuT tumors using tumor-level RNA-seq, histone ChIP-seq, and single-cell multiome profiling. PWK tumors exhibited broad transcriptional reprogramming characterized by reduced immune-associated expression and increased epithelial differentiation and neuronal-like programs. Chromatin remodeling was spatially redistributed rather than uniformly increased or decreased. PWK tumors showed expansion of H3K9me3- and H4K20me3-associated chromatin, while both marks were depleted across many expressed genes and concentrated within selected intergenic and repeat-rich territories, revealing coordinated redistribution of constitutive heterochromatin. Single-cell multiome profiling showed that broad epithelial representation remained similar between strains, yet specific malignant states were strongly strain biased. Integrated RNA and accessibility analyses identified concordant strain-selective regulatory programs and candidate loci, while trajectory analysis revealed distinct epithelial state transitions between PWK NeuT and BALB NeuT tumors. Together, these findings indicate that inherited genetic background reshapes transcriptional programs, heterochromatin organization, and malignant epithelial state trajectories in NeuT mammary tumors, providing a regulatory framework for understanding strain-dependent differences in tumor onset.

## INTRODUCTION

Cancer-driving mutations are interpreted within an inherited genome that can shape the regulatory response to oncogenic signaling. Well-known germline variations influence breast cancer susceptibility^1–10^. In addition, many common risk variants reside outside protein-coding sequence within distal regulatory elements and transcription-factor binding sites^11^. Yet association studies generally identify loci rather than the molecular consequences of inherited variation within an evolving tumor. How genetic background alters chromatin organization, transcriptional state, and the range of cellular programs accessible to cells carrying the same oncogenic driver therefore remains poorly understood.

Genetically controlled mouse models provide an opportunity to isolate this interaction because the initiating oncogene can be held constant while inherited background varies. The BALB/c background MMTV-NeuT model expresses constitutively active rat Neu/ErbB2 in mammary epithelium and develops timely spontaneous multifocal mammary tumors. Previous analysis of NeuT mice on genetically diverse backgrounds demonstrated that inherited variation strongly modifies tumor latency and growth^12^. Quantitative-trait loci associated with tumor onset mapped to chromosomes 1 and X, and genetic transfer experiments showed that the PWK background accelerates NeuT mammary tumor development ^12^. These observations establish a phenotype that is not explained by the initiating oncogene itself and raise a central question: what regulatory features of the PWK genome alter the response of mammary epithelial cells to NeuT and permit earlier tumor development?

PWK/PhJ is a wild-derived Mus musculus strain with extensive sequence divergence from classical laboratory strains. PWK and related wild-derived genomes contain abundant single-nucleotide and structural variation relative to standard laboratory references, providing a broad source of inherited regulatory diversity^13, 14^. A substantial fraction of structural variation among diverse mouse strains is associated with transposable elements, and such variation can alter local chromatin accessibility^15^. Strain-polymorphic endogenous retroviral insertions can themselves function as cis-regulatory elements, altering nearby H3K27ac and transcription^16^. Thus, inherited sequence differences can influence regulatory activity both through conventional cis-regulatory variants and through variation in repetitive genomic sequence. These features make PWK a particularly informative background for asking how inherited genomic architecture modifies an oncogene-driven tumor phenotype, while also requiring careful interpretation of reference-dependent mapping and repeat-associated signals.

Chromatin provides a plausible interface between inherited sequence and tumor cell state. H3K4me3 and H3K27ac mark active promoters and regulatory elements, whereas H3K9me3 and H4K20me3 are associated with constitutive heterochromatin, repetitive sequence, and large repressive genomic domains. Importantly, the biological consequence of these marks depends on their genomic distribution rather than total abundance alone. H3K9me3-enriched domains can restrict transitions between cellular states, whereas Suv39h-dependent H3K9me3 preferentially marks and represses intact LINE and endogenous retroviral elements^17, 18^. In a genetically divergent tumor background, redistribution of repressive chromatin among genes, intergenic regions, and repetitive elements could therefore alter which transcriptional programs remain accessible during tumor evolution.

Genetic background may also act by changing the cellular states selected under oncogenic pressure. Tumors driven by the same initiating lesion can contain distinct malignant epithelial programs even when their broad cellular composition is similar. Joint single-cell RNA and chromatin-accessibility profiling provides a means to resolve these differences by defining multimodal cell states and linking transcriptional programs to accessible regulatory elements and transcription-factor-associated motifs^19–21^. This distinction is particularly relevant to tumor latency: accelerated tumor formation need not require wholesale replacement of cellular compartments, but could instead reflect altered occupancy or accessibility of malignant epithelial states that are more compatible with NeuT-driven growth.

Here, we integrated tumor-level RNA-seq and ChIP-seq profiling of H3K4me3, H3K27ac, H3K9me3, and H4K20me3 with single-cell multiome analysis of PWK NeuT and BALB NeuT mammary tumors. We asked whether the PWK background associated with accelerated tumor onset is accompanied by coordinated changes in transcription, chromatin organization, repetitive-element-associated heterochromatin, and malignant epithelial state structure. We find that PWK tumors are characterized by reduced immune-associated transcriptional programs, extensive redistribution of H3K9me3 and H4K20me3 between gene-associated and repeat-rich genomic compartments, and marked reorganization of malignant epithelial regulatory states and trajectories. Together, these findings identify inherited genetic background as a determinant of the regulatory landscape through which a shared NeuT driver is expressed and provide a genomic framework for understanding accelerated tumor development in the PWK background.

## RESULTS

### Genetic background drives major transcriptional and chromatin differences in NeuT tumors

We compared PWK NeuT and BALB NeuT mammary tumors using matched bulk RNA-seq and ChIP-seq profiling of H3K4me3, H3K27ac, H3K9me3, and H4K20me3, together with single-cell multiome profiling (**Fig. 1A**). Bulk RNA-seq and ChIP-seq defined strain-associated transcriptional and chromatin differences, whereas the independent single-cell multiome cohort was used to determine how these differences were distributed across tumor cell states.

**Figure 1.**
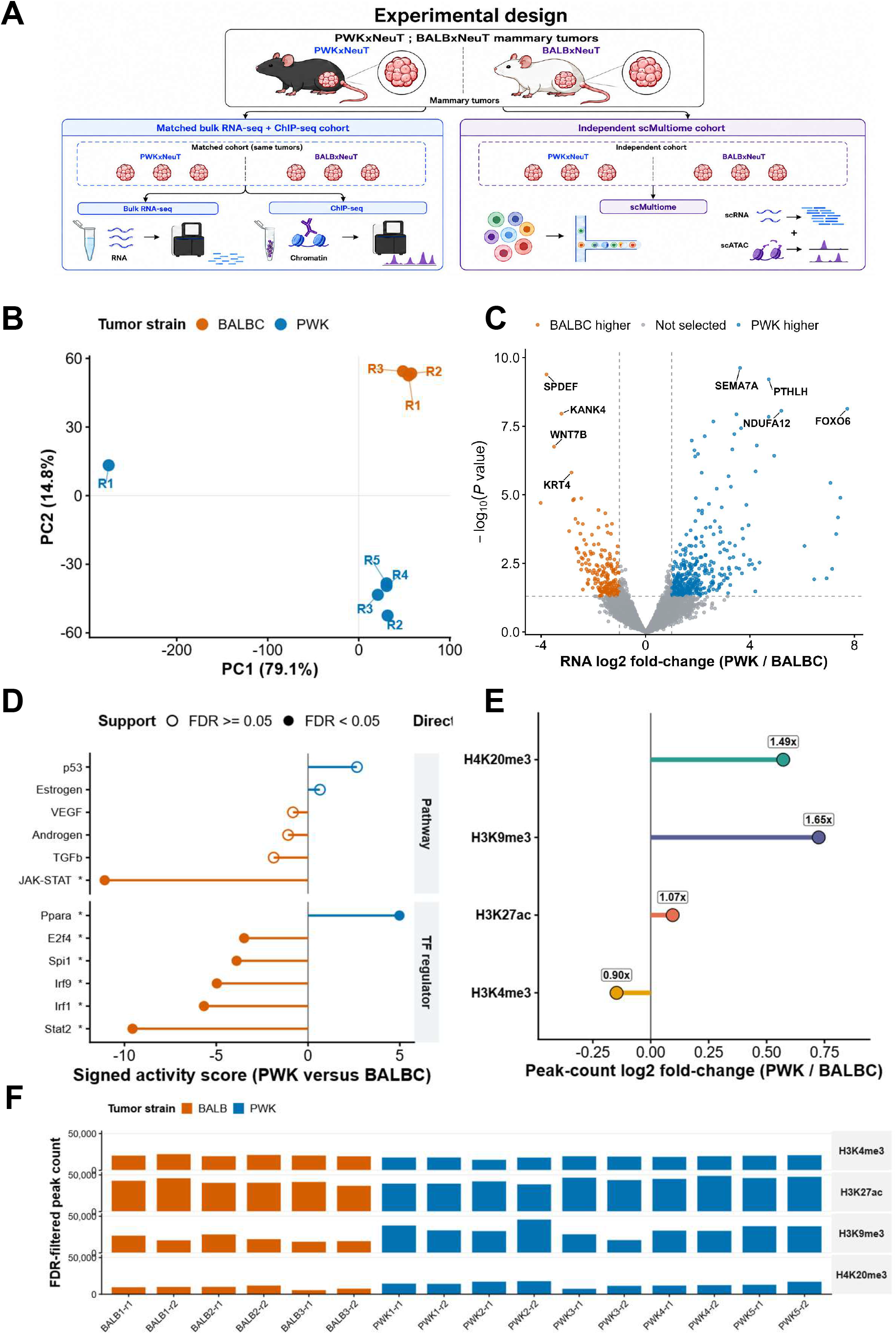
Genetic background defines the dominant transcriptome and histone-landscape axis in NeuT mammary tumors. (A) Experimental design integrating matched bulk RNA-seq and ChIP-seq with an independent single-cell multiome cohort. (B) Tumor-level RNA principal-component analysis showing separation by genetic background. (C) Bulk RNA differential-expression analysis. Positive log2 fold change indicates higher expression in PWK and negative values indicate higher expression in BALB/c. (D) Signed pathway and regulator activity scores. Positive values indicate higher inferred activity in PWK and negative values indicate higher activity in BALB/c; filled symbols indicate FDR < 0.05. (E) Strain-associated changes in ChIP-seq peak burden for H3K4me3, H3K27ac, H3K9me3, and H4K20me3, expressed as log2(PWK/BALB/c). (F) Filtered peak counts across individual ChIP-seq libraries.

Genetic background dominated the bulk transcriptomic landscape. The first principal component accounted for 79.1% of expression variance and separated PWK NeuT from BALB NeuT tumors, whereas the second principal component accounted for substantially less variation (**Fig. 1B**). Differential-expression analysis identified 561 strain-associated genes, with 378 increased in PWK and 183 increased in BALB/c tumors (**Fig. 1C**). These differences were broadly distributed across the transcriptome, indicating that genetic background altered the transcriptional state of NeuT tumors rather than a restricted set of individual genes. Regulatory and pathway-level analyses further distinguished the strains, with stronger JAK-STAT and STAT2/IRF-associated activity in BALB NeuT tumors and increased PPARA-associated activity in PWK NeuT tumors (**Fig. 1D**). Together, these findings define a prominent strain-dependent transcriptional axis in which PWK tumors show reduced immune-associated regulatory activity and a distinct metabolic transcriptional state.

Genetic background also produced marked differences in the histone landscape (**Fig. 1E-F**). The largest changes involved the constitutive heterochromatin-associated marks H3K9me3 and H4K20me3, which showed substantially greater peak burden in PWK tumors, whereas changes in H3K27ac and H3K4me3 were comparatively modest. These differences suggest that the PWK phenotype is associated not simply with altered transcription, but with large-scale reorganization of repressive chromatin.

Together, these data establish genetic background as the principal source of molecular variation among NeuT tumors in this cohort and identify two prominent features of the PWK state: broad attenuation of immune-associated transcriptional programs and disproportionate remodeling of H3K9me3- and H4K20me3-associated chromatin.

### PWK tumors redistribute repressive chromatin away from gene-associated regions

Expression-stratified metagene analysis showed that the greater overall H3K9me3 and H4K20me3 peak burden in PWK tumors was not accompanied by increased signal across gene-associated regions (**Fig. 2A-C**). Genes were grouped according to underlying expression level, and histone-modification signal was examined across promoters, gene bodies, and downstream regions. H3K9me3 signal was lower in PWK across these gene-centered regions in all expression strata, with the largest reduction around highly expressed genes. Among highly expressed genes, H3K9me3 signal was reduced by approximately 20 to 25% in PWK across promoter-proximal, gene-body, and downstream regions. H4K20me3 showed a similar but more modest reduction, with signal decreased by approximately 10 to 12% across highly expressed gene-associated regions. In contrast, H3K4me3 was increased near promoters in PWK, including an approximately 17 to 19% increase in the highest and middle expression quartiles, whereas H3K27ac differences were smaller and more context dependent.

**Figure 2.**
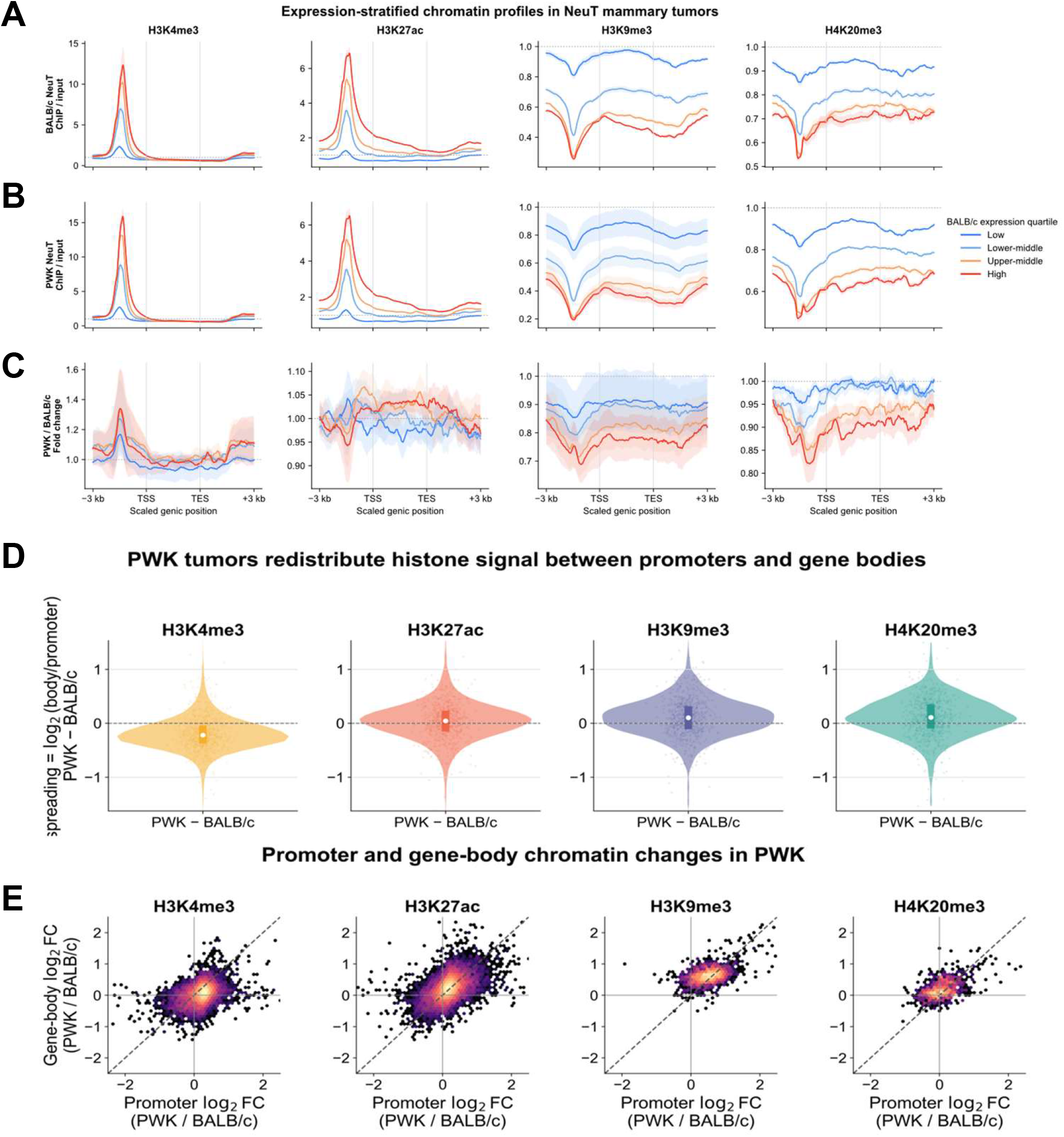
PWK tumors redistribute histone signal between promoters, gene bodies, and non-genic compartments. (A-B) Expression-stratified metagene profiles for H3K4me3, H3K27ac, H3K9me3, and H4K20me3 in BALB/c (A) and PWK (B) tumors. Genes were grouped by underlying expression level and profiled from 3 kb upstream of the transcription start site through the scaled gene body to 3 kb downstream of the transcription end site. Lines show strain summaries and ribbons show between-tumor variation. (C) PWK-to-BALB/c signal ratios across the same intervals; values below 1 indicate lower signal in PWK. (D) Per-gene change in the log2 gene-body-to-promoter signal ratio between PWK and BALB/c. (E) Relationship between promoter and gene-body log2 fold changes for each histone modification. The diagonal denotes equal promoter and gene-body effects.

Promoter-to-gene-body comparisons further showed that each histone modification was redistributed in a distinct manner (**Fig. 2D-E**). H3K4me3 shifted relatively toward promoters in PWK tumors, whereas H3K9me3 and H4K20me3 shifted relatively toward gene bodies. Thus, the strain effect involved not only differences in overall histone-mark abundance, but also systematic redistribution of signal across genic compartments that was not apparent from peak counts alone.

Together, these analyses reveal a marked spatial redistribution of repressive chromatin in PWK tumors. Despite greater overall H3K9me3- and H4K20me3 peak burden, many gene-associated regions show reduced signal for both marks, indicating that the PWK chromatin phenotype reflects redistribution across genomic compartments rather than a uniform increase in repressive histone modifications.

### Combinatorial chromatin states reveal selective redistribution of repressive chromatin

To determine how the increased H3K9me3 and H4K20me3 peak burden in PWK tumors was distributed across the genome, we learned a 16-state ChromHMM^22^ model from the four histone marks without assigning state identities a priori (**Fig. 3A**). PWK tumors did not show a generalized increase across all repressive states. Instead, the largest gains were confined to an H3K9me3-dominant state and a dual H3K9me3-H4K20me3 state, whereas other repressive states changed little or contracted (**Fig. 3B**). This pattern indicates selective expansion and contraction of specific combinatorial chromatin states rather than a uniform genome-wide increase in repressive chromatin.

**Figure 3.**
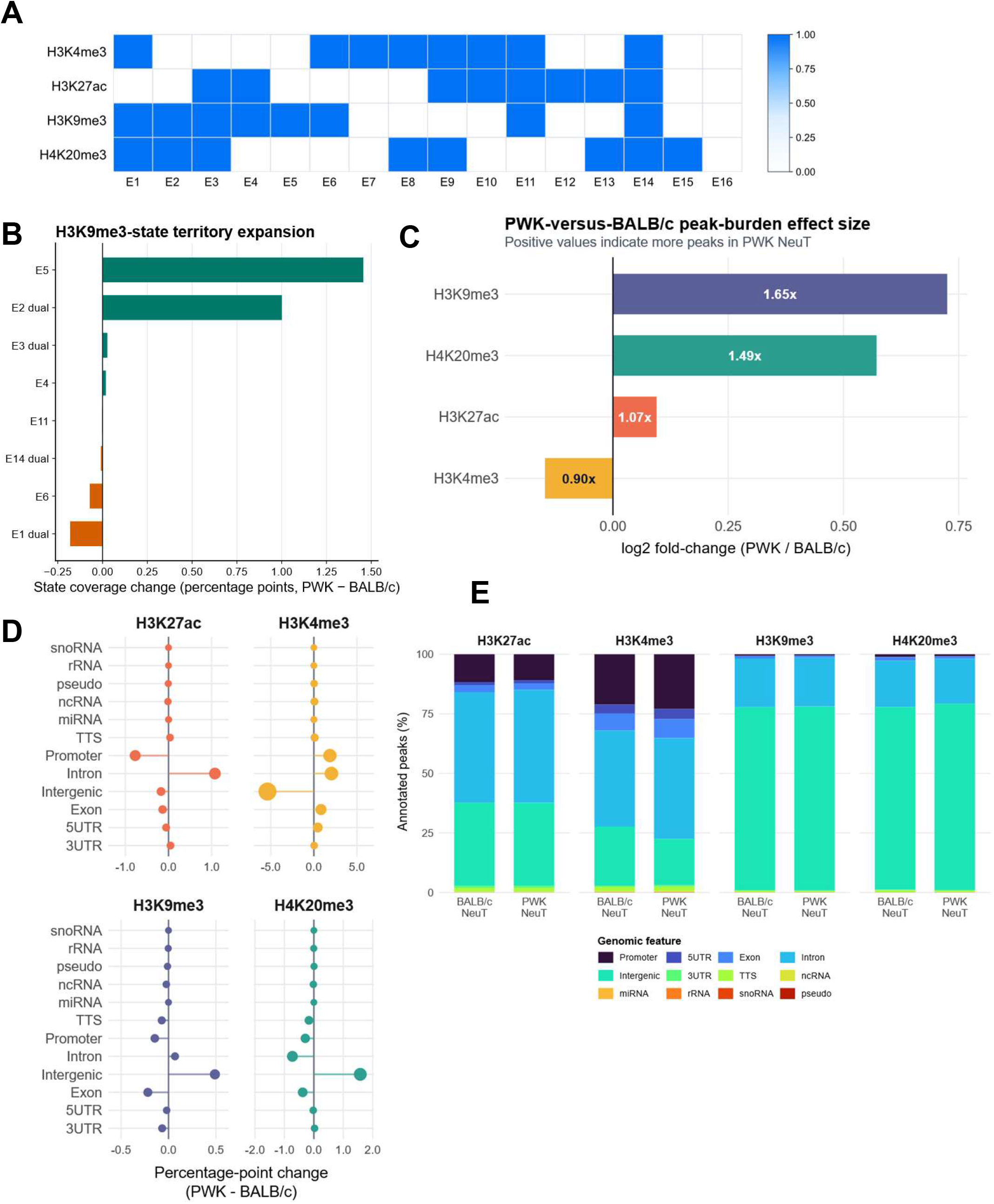
Combinatorial chromatin states resolve selective expansion of intergenic repressive territories in PWK tumors. (A) Emission matrix for the 16-state ChromHMM model learned from H3K4me3, H3K27ac, H3K9me3, and H4K20me3. (B) PWK-minus-BALB/c changes in genomic coverage for H3K9me3-containing and dual H3K9me3-H4K20me3 chromatin states. (C) Strain-associated peak-burden changes for the four histone modifications, expressed as log2(PWK/BALB/c). (D) PWK-minus-BALB/c changes in the genomic-feature composition of histone-enriched regions. Positive values indicate greater representation in PWK. (E) Mean genomic-feature composition of histone-enriched regions for each mark and strain after summarization within biological tumors.

Genomic annotation analysis further showed that strain-associated chromatin remodeling was spatially selective (**Fig. 3D-E**). In PWK tumors, H3K4me3 became relatively more concentrated at promoters and less represented in intergenic regions. In contrast, H3K9me3 and H4K20me3 shifted toward intergenic sequence, with corresponding reductions across several promoter and other gene-associated annotations. Heatmaps centered on strain-specific enriched regions showed that these patterns were consistently represented across biological tumors rather than arising solely from aggregate peak calls (**Fig. S1**).

Broad-domain analysis further distinguished the behavior of H3K9me3 and H4K20me3. H4K20me3 domain coverage was greater in PWK tumors, whereas broad H3K9me3 coverage was similar between strains. Regions jointly occupied by H3K9me3 and H4K20me3 were also more extensive in PWK, with greater overlap between the two marks. Thus, the increased H3K9me3 peak burden in PWK does not reflect uniform expansion of broad H3K9me3 domains. Instead, the PWK chromatin landscape is characterized by selective expansion of specific H3K9me3-containing chromatin states, broader H4K20me3 coverage, increased overlap between the two repressive marks, and redistribution toward intergenic genomic compartments.

### Repeat-rich sequence anchors coordinated heterochromatin reorganization

Repeat-centric analysis localized much of the strain-associated repressive chromatin remodeling to selected ERV, LTR, LINE, and satellite territories (**Fig. 4A**). ERV1 and ERVK families showed increased H3K9me3- and H4K20me3-associated enrichment in PWK tumors, whereas satellite sequences displayed a more mark-selective response. Consistent with coordinated remodeling of the two repressive marks, PWK-minus-BALB/c changes in H3K9me3 and H4K20me3 were strongly correlated across fixed genomic windows (**Fig. 4B**). Repeat-territory enrichment and peak-centered metaprofiles further showed increased PWK signal across selected repressive regions (**Fig. 4C-D**).

**Figure 4.**
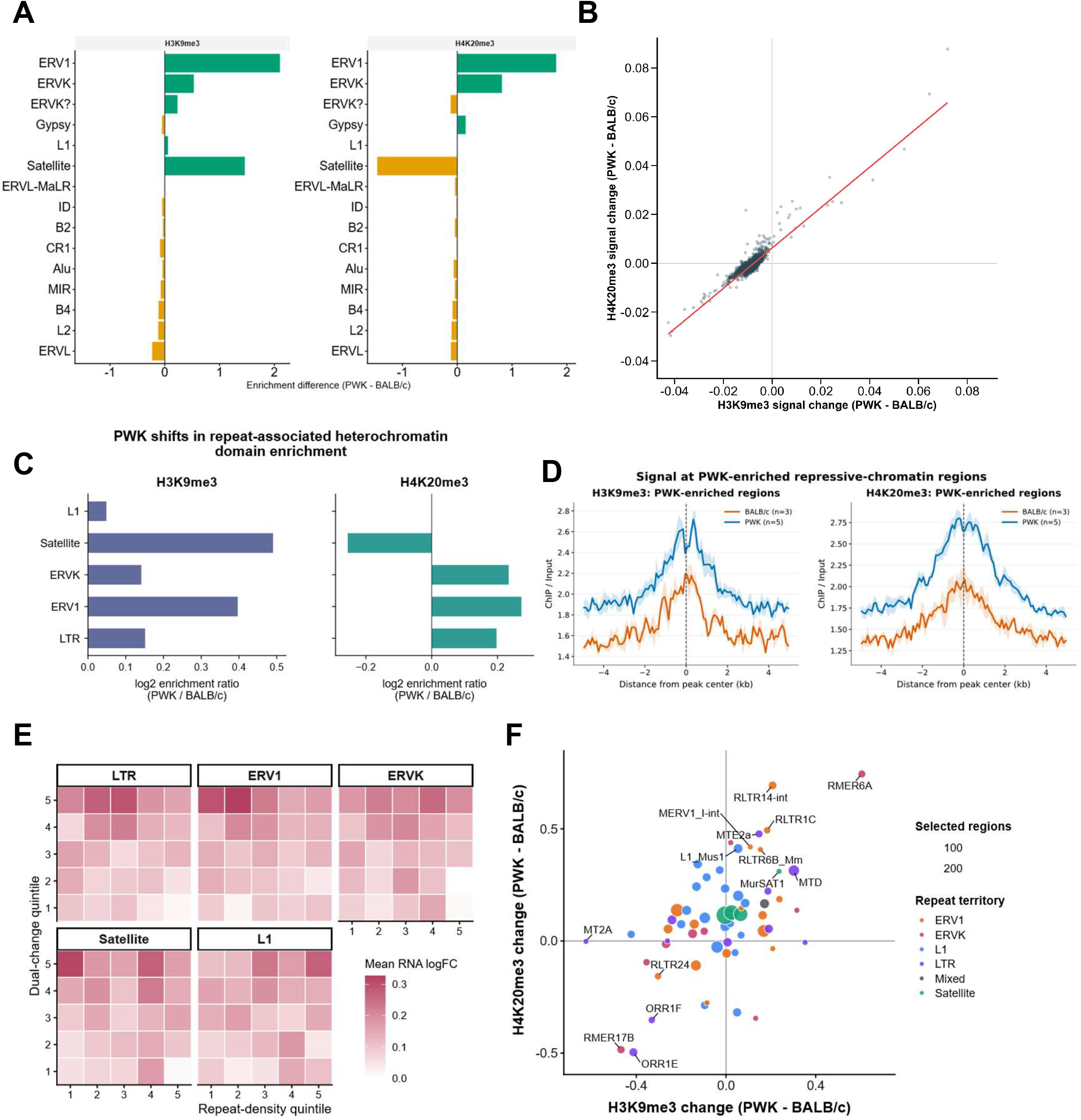
Repeat-rich sequence anchors coordinated H3K9me3-H4K20me3 reorganization. (A) PWK-minus-BALB/c differences in repeat-family enrichment within H3K9me3- and H4K20me3-associated regions. Positive values indicate greater PWK enrichment. (B) Relationship between PWK-minus-BALB/c H3K9me3 and H4K20me3 signal changes across fixed 1-Mb genomic windows. Changes in the two repressive marks were positively correlated across the genome (Spearman ρ = 0.75). (C) PWK-to-BALB/c enrichment ratios for repeat territories within strain-associated repressive domains. (D) Mean ChIP/input signal centered on PWK-enriched repressive regions across biological tumors. (E) Mean bulk RNA log2 fold change across repeat-density and dual-mark-change quintiles for major repeat territories. (F) H3K9me3- and H4K20me3-effect landscape for selected H3K9me3-enriched repeat subfamilies. Point color denotes repeat territory and point size denotes the number of selected regions.

The relationship between repeat-associated heterochromatin and transcription was not monotonic (**Fig. 4E**). Genes located within several repeat-dense genomic neighborhoods that gained both repressive marks nevertheless showed positive mean RNA effects in PWK. This pattern does not imply that H3K9me3 or H4K20me3 promotes transcription. Rather, it indicates that repeat-associated chromatin remodeling and nearby transcriptional output can vary independently across genomic contexts, consistent with additional contributions from local regulatory architecture and tumor-cell state composition.

Analysis of H3K9me3-enriched repeat regions further resolved coordinated and mark-selective changes that were obscured by broader repeat-class annotations (**Fig. 4F; Fig. S2**). Within this H3K9me3-marked subset, several repeat subfamilies, including RMER6A, RLTR14-int, RLTR1C, MTE2a, MERV1_I-int, and selected L1-associated elements, showed higher H3K9me3 and/or H4K20me3 signal in PWK, whereas others, including RMER17B, ORR1E, ORR1F, MT2A, and RLTR24, were shifted toward BALB/c. These findings indicate that strain-associated heterochromatin remodeling is concentrated within specific H3K9me3-marked repeat subfamilies rather than distributed uniformly across repetitive sequence, supporting a broader model of selective heterochromatin redistribution in PWK tumors.

### Genetic background redistributes malignant epithelial states without altering epithelial dominance

Single-cell multiome profiling integrating gene expression and chromatin accessibility measurements (GEX + ATAC) resolved 34 weighted-nearest-neighbor clusters spanning malignant epithelial, immune, stromal, vascular, and minor cell populations (**Fig. 5A-C**). Broad epithelial representation was similar between PWK NeuT and BALB NeuT tumors, indicating that genetic background did not substantially alter the overall epithelial dominance of the tumors. Instead, the major strain effect was a redistribution among malignant epithelial states.

**Figure 5.**
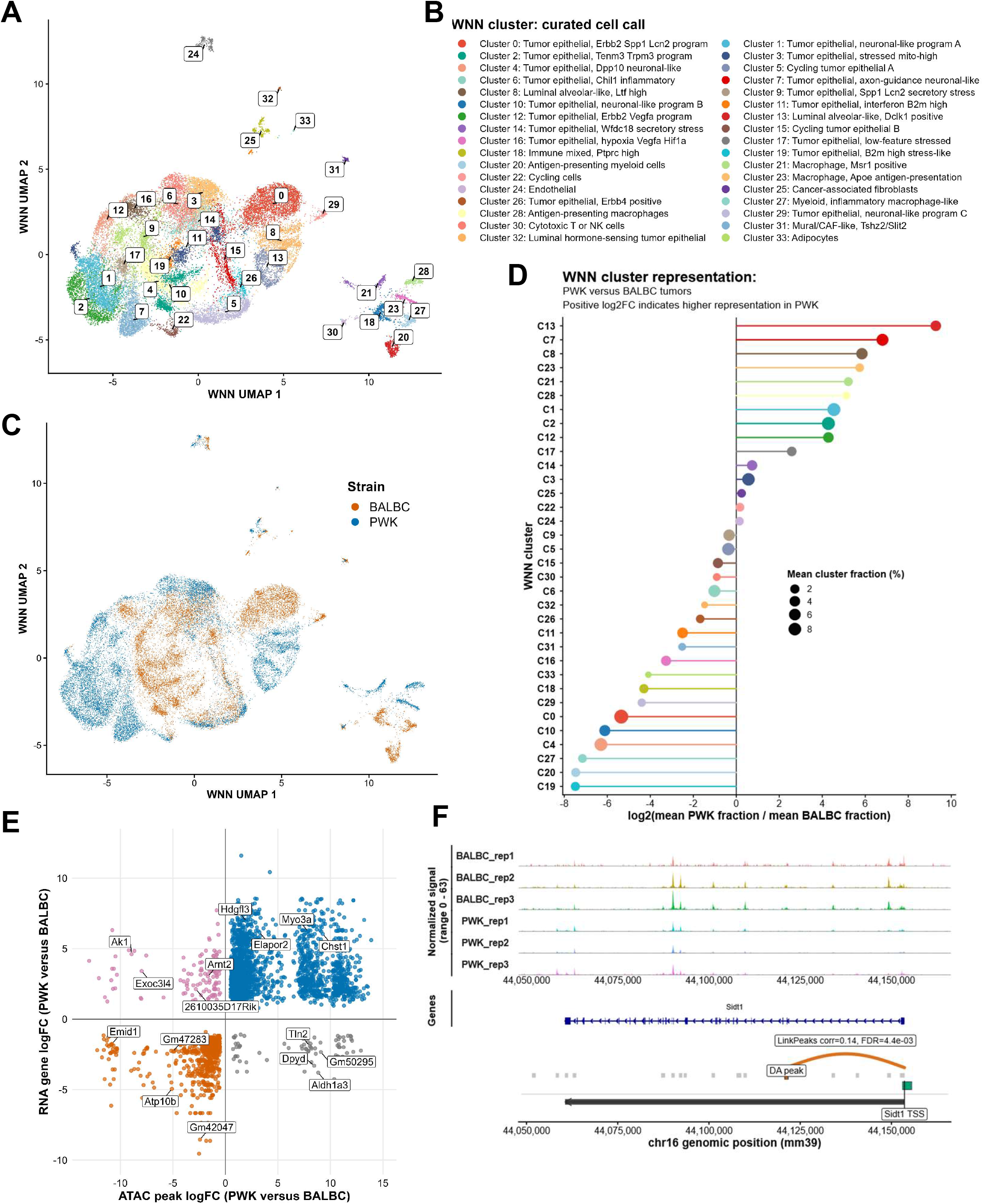
Single-cell multiome profiling resolves strain-biased malignant states and concordant cis-regulatory changes. (A) WNN UMAP of 32,304 cells profiled by joint gene expression and chromatin accessibility, colored by multimodal cluster. (B) Curated cluster-to-cell-state annotations. (C) WNN UMAP colored by strain. (D) Relative representation of each WNN cluster in PWK versus BALB/c tumors, expressed as log2(mean PWK fraction/mean BALB/c fraction); point size denotes mean cluster abundance. (E) Integrated tumor-level differential ATAC and linked RNA effects within sufficiently represented cell states. Positive values indicate higher signal in PWK. (F) Representative BALB/c-high locus showing normalized ATAC coverage across biological tumors, gene annotation, a differential-accessibility peak, and a peak-to-gene link.

Several epithelial states were strain biased (**Fig. 5D; Fig. S3**). Cluster-associated RNA marker patterns supported the curated epithelial, immune, stromal, vascular, and minor cell-state assignments across the WNN atlas (**Fig. S4**). PWK tumors were enriched for neuronal-like, axon-guidance, Tenm3/Trpm3, Ltf-high luminal-alveolar, and Dclk1-positive luminal-alveolar states, whereas BALB NeuT tumors preferentially occupied ErbB2/Spp1/Lcn2, Dpp10 neuronal-like, interferon/B2m-high, and hypoxic epithelial states. Some of these differences were nearly strain restricted, indicating that genetic background strongly reshapes the distribution of malignant epithelial programs under a shared NeuT driver.

Erbb2-expressing epithelial states were represented in both strains but differed in their accompanying transcriptional programs. A PWK-enriched Erbb2/Vegfa state contrasted with a BALB/c-enriched Erbb2/Spp1/Lcn2 state, indicating that genetic background is associated with distinct Erbb2-expressing epithelial programs. Myeloid states also differed between strains. BALB NeuT tumors contained B2m-high, antigen-presentation, and inflammatory myeloid states, whereas PWK NeuT tumors showed increased representation of macrophage populations, including Apoe-positive antigen-presenting and Msr1-positive states. Together, these findings indicate that the PWK background primarily alters the spectrum of malignant epithelial and immune-associated states rather than the broad cellular composition of NeuT tumors.

### Concordant RNA-accessibility changes identify strain-biased regulatory loci

We next asked whether strain-biased tumor states were accompanied by coordinated changes in gene expression and chromatin accessibility. Tumor-level pseudobulk analysis within sufficiently represented states identified 3,071 differential ATAC peak-gene pairs involving 1,828 accessible regions and 603 genes (**Fig. 5E**). Most pairs showed concordant directionality between accessibility and RNA, with PWK-high accessibility preferentially linked to PWK-high expression and the reciprocal pattern observed in BALB NeuT tumors. Because these comparisons were performed within annotated cell states, the concordance reflects state-resolved regulatory differences rather than simply differences in state abundance.

Concordant PWK-high loci included Sema7a, Tcerg1l, Frem1, Elapor2, Myo3a, Chst1, Enpp2, Asic2, and Robo2, whereas BALB/c-high loci included Sidt1, Atp10b, Kank4, Vipr1, Epsti1, Pik3cg, Slc38a1, Zfp423, and Lcp1. At the Sidt1 locus, accessibility was consistently greater in BALB NeuT tumors, accompanied by higher Sidt1 expression and a linked differential accessibility peak (**Fig. 5F**). Reciprocal PWK-high patterns were observed at Sema7a, Frem1, Tcerg1l, and Elapor2, where RNA direction, differential accessibility, and locus-level coverage converged across tumors (**Fig. S5**).

Cross-platform integration further identified genes supported by bulk RNA-seq, state-resolved single-cell expression, chromatin accessibility, and active histone-mark evidence (**Fig. S6A**). Tcerg1l, Chst1, Fetub, Foxd2os, Samd5, Dock10, Ncam1, Enpp2, Chst2, Pde10a, Moxd1, Atp10a, and Frem1 were among the PWK-high candidates supported across multiple genomic layers. Together, these analyses define a focused set of strain-responsive regulatory loci in which transcriptional and chromatin differences converge. Agreement across independent genomic modalities prioritizes candidates for subsequent allele-aware and functional analysis.

### State-resolved motif activity defines regulatory circuits embedded in strain-biased populations

Integrated analysis of transcription-factor expression and chromVAR^21^ motif activity revealed distinct regulatory programs across the 34 WNN states (**Fig. 6A**). These patterns were strongly state specific rather than reflecting uniform strain-wide changes. For example, the cycling epithelial C5 state showed concordant SOX6- and PLAGL1-associated expression and motif signals, whereas the Spp1/Lcn2 secretory-stress state was characterized by FOXO1-associated activity and the hypoxic epithelial C16 state by BACH1::MAFK-, CREB5-, and ARNT-associated motif accessibility. Distinct programs were also evident in non-epithelial populations, including SMAD2/3-, MAFB-, and SOX11-associated signals in endothelial cells and RUNX1-, BACH1-related, and ARNTL-associated signals in cancer-associated fibroblasts.

**Figure 6.**
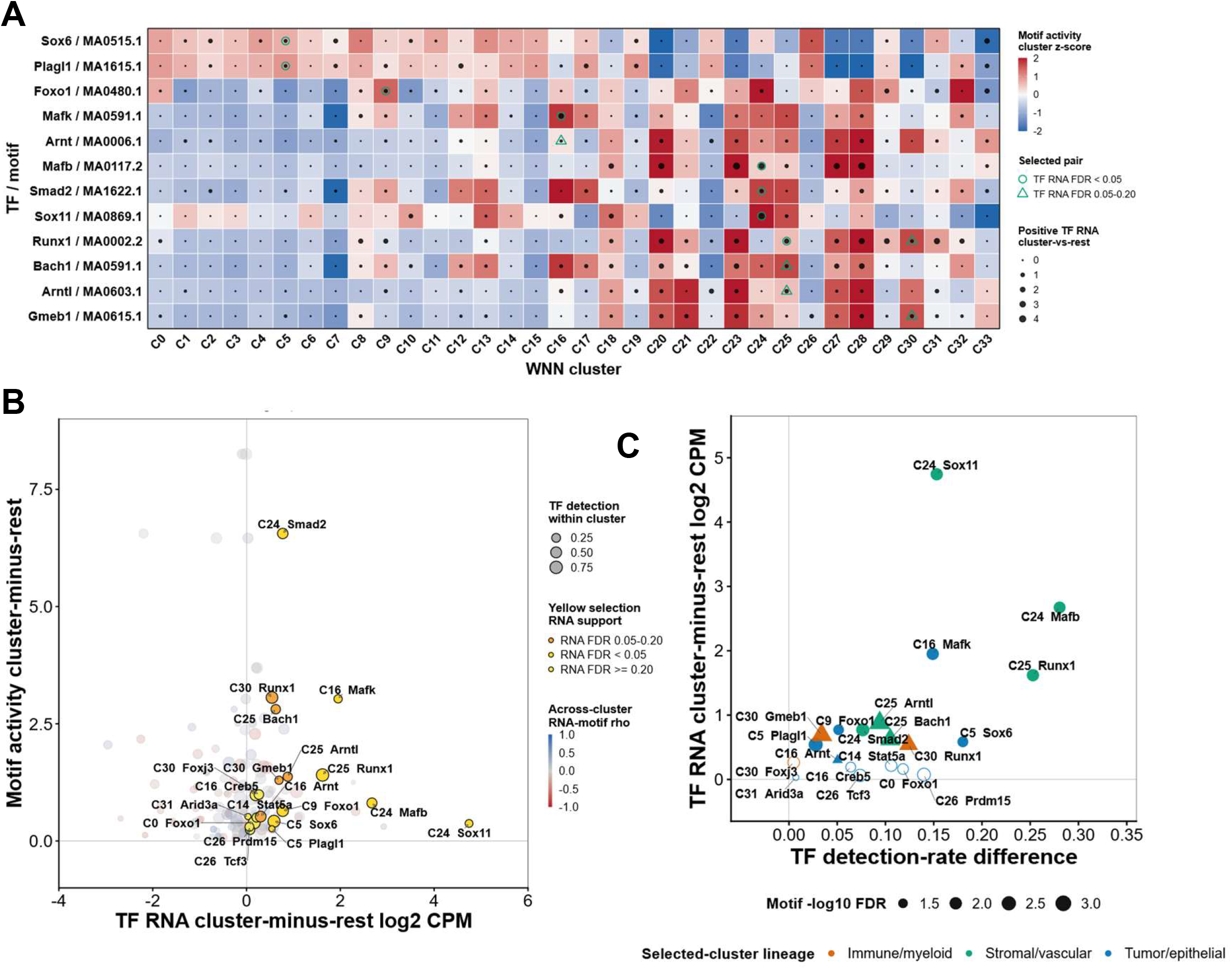
Replicate-supported TF RNA and motif activity define regulatory circuits within distinct tumor states. (A) Matrix of transcription-factor RNA expression and chromVAR motif activity across 34 WNN states. Tile color denotes normalized motif activity, and overlaid symbols denote transcription-factor RNA specificity and statistical support as defined in the panel. (B) Relationship between selected-state transcription-factor RNA and motif activity. Point size denotes transcription-factor detection within the selected state, and color denotes cross-state RNA-motif correlation. (C) Relationship between transcription-factor detection-rate difference and selected-state RNA expression. Point size denotes motif significance, color denotes cell-state lineage, and symbols denote RNA statistical support. Motif activity reflects accessibility at motif-containing regions and does not establish direct transcription-factor binding.

For several factors, changes in motif activity were accompanied by concordant changes in expression of the corresponding transcription factor, including SOX6 in C5, BACH1::MAFK-associated activity with Mafk expression in C16, SMAD2/3-associated activity with Smad2 expression in C24, and RUNX1 in C25 (**Fig. 6A**). Other motifs showed strong state selectivity without corresponding changes in cognate RNA, consistent with motif accessibility and transcription-factor abundance capturing distinct regulatory layers. Across states, TF expression and motif accessibility provided complementary rather than redundant information, with concordance restricted to a subset of regulatory programs (**Fig. 6B-C**).

Together, these findings show that strain-biased tumor states are associated with distinct combinations of transcription-factor expression and motif-bearing chromatin accessibility. Genetic background can therefore alter the tumor regulatory landscape in part by changing the representation of cellular states that carry different regulatory programs, without requiring uniform activation of a single transcription factor across the tumor. Because chromVAR measures accessibility at motif-containing regions rather than direct transcription-factor occupancy, these associations identify candidate regulators for subsequent functional and binding-based validation.

### Genetic background reshapes epithelial-state trajectories

We modeled malignant epithelial-state organization using independently fitted axon-guidance epithelial and stress-secretory trajectories (**Fig. 7A**). The axon-guidance epithelial trajectory was fitted after restricting the analysis to malignant epithelial states. The inferred topologies were evaluated across RNA, ATAC, joint RNA-ATAC, and WNN representations and for stability across biological replicates. Recurrently recovered relationships connected the Tenm3/Trpm3, axon-guidance, and related epithelial states, whereas the stress-secretory trajectory connected cycling, interferon/B2m, secretory-stress, hypoxic, and other epithelial states. These analyses support organization of the malignant epithelial compartment into multiple transcriptionally related paths rather than a single linear continuum. Because pseudotime orders transcriptionally related states, the inferred trajectories should not be interpreted as direct evidence of developmental ancestry.

**Figure 7.**
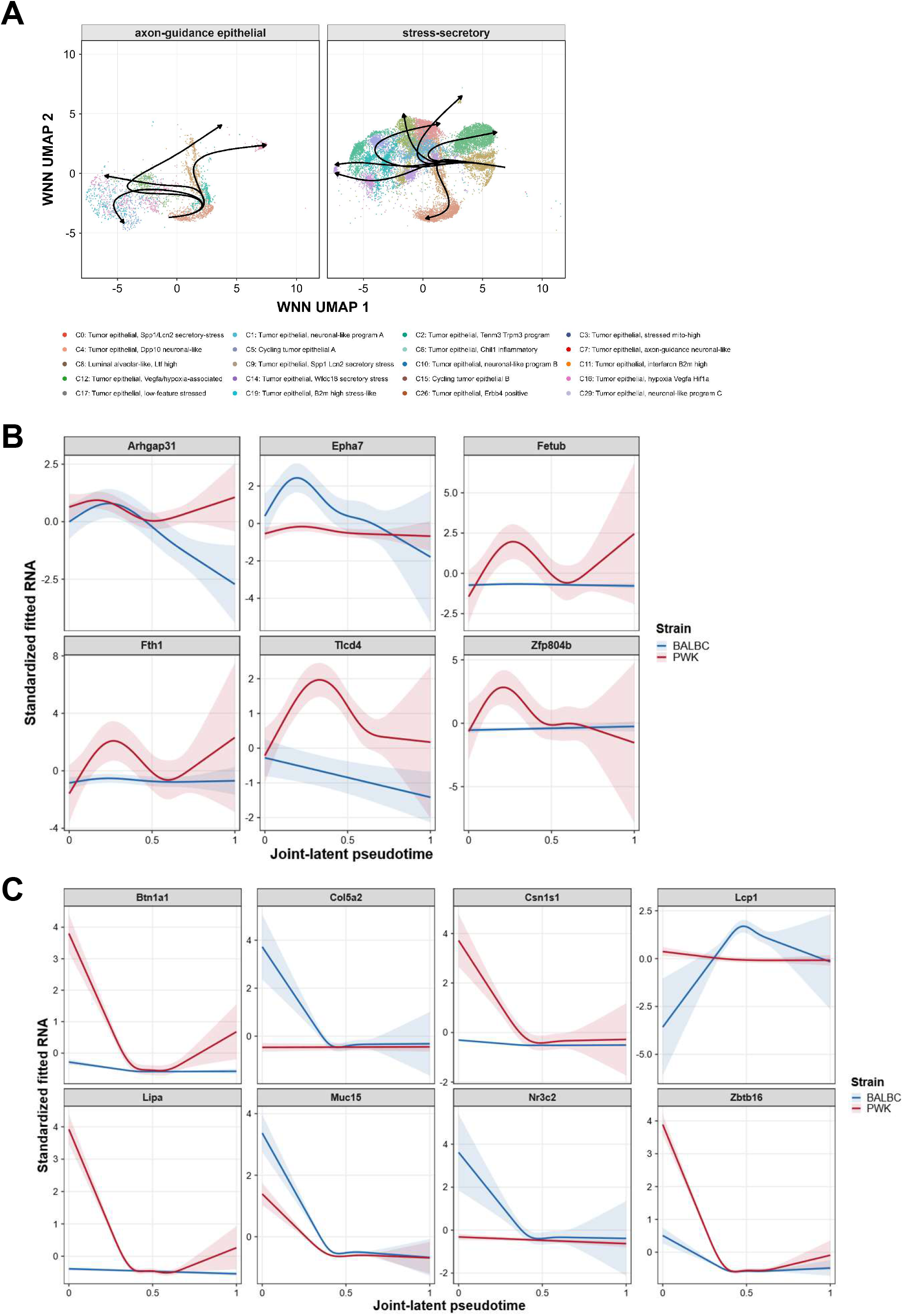
Genetic background redirects malignant epithelial trajectories. (A) Independently fitted axon-guidance epithelial and stress-secretory Slingshot trajectories displayed on the shared WNN embedding. Arrows indicate pseudotemporal orientation. Trajectory topology was evaluated across RNA, ATAC, joint RNA-ATAC, and WNN representations and for stability across biological replicates. (B) Strain-dependent transcriptional dynamics along the axon-guidance epithelial trajectory. (C) Strain-dependent transcriptional dynamics along the stress-secretory trajectory. Curves show fitted standardized RNA values, and shaded regions indicate uncertainty. Pseudotime represents an inferred ordering of transcriptionally related states and does not establish developmental ancestry.

Along the axon-guidance epithelial trajectory, six genes showed robust strain-dependent expression dynamics based on prespecified statistical and epithelial-specificity criteria (**Fig. 7B**). Arhgap31, Epha7, Fetub, Fth1, and Tlcd4 showed higher fitted expression in PWK than in BALB/c across portions of the trajectory, whereas Zfp804b showed lower fitted expression in PWK at later pseudotime. These results identify strain-dependent transcriptional divergence along the axon-guidance epithelial trajectory.

The independently fitted stress-secretory trajectory showed a distinct set of strain-dependent transcriptional dynamics (**Fig. 7C**). Btn1a1, Csn1s1, Lipa, and Zbtb16 showed higher fitted expression in PWK, most prominently at early pseudotime, whereas Col5a2, Lcp1, Muc15, and Nr3c2 showed higher fitted expression in BALB/c across portions of the trajectory. Several differences diminished or changed magnitude across pseudotime, indicating dynamic rather than constant strain effects. Together with the strain-biased epithelial states identified in the WNN atlas, these results indicate that PWK and BALB/c genetic backgrounds are associated with distinct transcriptional trajectories within a shared malignant epithelial-state space.

### Integrated immune and chromatin programs are associated with accelerated PWK tumor onset

We next asked whether the molecular features of established PWK NeuT tumors were consistent with the previously observed accelerated-onset phenotype. Genes reduced in PWK were strongly enriched for antigen processing and presentation, interferon response, cell killing, cytokine-response, and related immune programs (**Fig. S6B**). This transcriptional pattern was consistent with reduced JAK-STAT and STAT2/IRF-associated activity and with the lower representation of interferon/B2m-high epithelial and antigen-presenting myeloid states in PWK tumors.

These immune-associated differences occurred alongside extensive chromatin remodeling. PWK tumors showed reduced H3K9me3 and H4K20me3 signal across many gene-associated regions, increased promoter-proximal H3K4me3, selective expansion of H3K9me3- and H4K20me3-chromatin in intergenic and repeat-rich compartments, and redistribution toward distinct neuronal-like and luminal-alveolar epithelial states. These findings suggest that the accelerated PWK phenotype is associated with coordinated changes in both immune-associated transcriptional programs and the regulatory landscape of malignant epithelial cells.

Together, these findings are consistent with a model in which reduced interferon-linked immune restraint and altered chromatin-state organization accompany a tumor environment more permissive to NeuT-driven epithelial state transitions. In this model, diminished immune-associated programs could reduce constraints on emerging tumor cells, while redistribution of gene-associated and repeat-associated chromatin could alter the range of epithelial transcriptional states accessible during tumor development. The concentration of H3K9me3 and H4K20me3 within selected repeat-rich regions further indicates that this regulatory remodeling is spatially organized rather than reflecting generalized loss of repressive chromatin.

Because all genomic assays were performed on established tumors, these data do not establish whether immune attenuation or chromatin remodeling precedes accelerated tumor formation. Instead, they define a coherent molecular framework linking the PWK genetic background to the immune, chromatin, and epithelial-state features of tumors that arise with shorter latency, and generate specific predictions that can be tested in premalignant mammary epithelium.

## DISCUSSION

This study links an inherited modifier of mammary tumor latency to coordinated changes in transcription, heterochromatin organization, malignant epithelial state structure and reduced immune surveillance. Although PWK NeuT and BALB NeuT tumors retained similar broad epithelial composition, they differed in immune-associated transcription, active and repressive histone modifications, repeat-associated chromatin, single-cell accessibility, regulatory-state abundance, and epithelial trajectories. The central finding is therefore not a generalized increase or decrease in chromatin activity. Instead, the PWK background redistributes chromatin across gene-associated and repeat-rich genomic compartments while altering the malignant epithelial states that emerge under a shared NeuT driver.

One prominent feature of the PWK phenotype is attenuation of interferon-associated and immune-regulatory programs. PWK tumors showed reduced JAK-STAT and STAT2/IRF-associated activity, depletion of antiviral, interferon-beta, antigen-presentation, cytokine-response, and cell-killing programs, reduced representation of interferon/B2m-high epithelial states, and fewer antigen-presenting or inflammatory myeloid states. This pattern is particularly relevant to the accelerated-onset phenotype because disruption of type I interferon signaling in NeuT mice accelerates mammary tumor development and increases tumor multiplicity and tumor-initiating cell abundance^23^. These observations do not indicate that PWK tumors phenocopy IFNAR1 loss or that a specific PWK allele disrupts this pathway. Rather, they place the reduced interferon-associated state observed here within a biological framework in which diminished type I interferon signaling can facilitate Neu-driven tumor development. The modest change in total immune-cell representation and skewing toward tumor-associated macrophages clusters further suggests that the PWK phenotype reflects changes in both immune-state composition and tumor-cell-associated cytokine programs rather than wholesale loss of the immune compartment.

A second major feature was altered malignant epithelial-state organization. PWK tumors were enriched for Tenm3/Trpm3, axon-guidance, Ltf-high luminal-alveolar, and Dclk1-positive states, whereas BALBxNeuT tumors were enriched for ErbB2/Spp1/Lcn2, Dpp10 neuronal-like, interferon-stress, and hypoxic states. The ErbB2-associated state warrants a technical clarification: because the mouse GRCm39 reference lacked a separate rat NeuT sequence, Erbb2 counts could not distinguish endogenous mouse transcripts from any NeuT-derived reads that mapped to the mouse locus. These differences were supported by state abundance, RNA expression, chromatin accessibility, regulatory peak-gene associations, motif accessibility, and trajectory analysis. Notably, neuronal-like programs were not uniformly associated with either genetic background. Distinct neuronal-like states were present in both PWK and BALBxNeuT tumors, indicating that inherited variation reshapes a broader epithelial plasticity landscape rather than simply activating or suppressing a single neuronal-like program.

The heterochromatin data provide a genomic framework for understanding how such state differences could arise. PWK tumors contained more H3K9me3- and H4K20me3-enriched regions overall, yet both marks were reduced across many gene-associated intervals. ChromHMM, genomic annotation, and broad-domain analyses further showed selective expansion of H3K9me3-containing and dual H3K9me3-H4K20me3 states, increased intergenic representation of both repressive marks, broader H4K20me3 coverage, and greater co-occupancy of H3K9me3 and H4K20me3. Thus, the PWK phenotype is better described as redistribution of constitutive heterochromatin than as generalized heterochromatin gain. Because H3K9me3-rich domains can constrain cell-state transitions and H3K9me3-dependent pathways repress selected intact retrotransposons^17, 18^, redistribution of these marks could alter the genomic regions that remain accessible during tumor evolution. The present data support this model.

Repeat-rich sequence represents an especially informative component of this redistribution. H3K9me3- and H4K20me3-associated changes were concentrated within selected ERV, LTR, LINE, satellite, and H3K9me3-enriched repeat subfamilies rather than uniformly across repetitive sequence. This observation is notable in PWK because wild-derived mouse genomes contain extensive sequence and structural variation relative to classical laboratory strains. Polymorphic endogenous retroviruses can generate strain-specific regulatory activity, and transposable-element-associated structural variants can alter nearby chromatin organization^15, 16^. The repeat-associated signals observed here may therefore reflect differences in the genomic substrate on which heterochromatin is assembled, rather than epigenetic remodeling of an identical sequence background. This interpretation also defines an important technical boundary: strain-specific repeat content and mappability must be separated from true chromatin-state differences using strain-aware or allele-aware genomic references.

Integration of RNA and chromatin accessibility further narrowed the strain-associated regulatory landscape to specific loci. Differential accessibility and expression were concordant for most peak-gene pairs, and loci including Sema7a, Tcerg1l, Frem1, Elapor2, Chst1, and Enpp2 were supported across multiple genomic layers. These loci should be viewed as prioritized regulatory candidates for tumor-latency genes.

The broader implication is that inherited cancer susceptibility may act not only through isolated risk genes but through genome-wide effects on regulatory-state accessibility. Many breast cancer risk variants occur in noncoding regulatory sequence^11, 24^, where their consequences are expected to depend on cell type, chromatin context, and oncogenic state. The PWK model illustrates how inherited variation can coincide with changes in immune-associated programs, repeat-associated heterochromatin, regulatory accessibility, and malignant epithelial state selection despite a common initiating oncogene. This framework suggests that the phenotypic effect of inherited variation may emerge from the combined organization of regulatory elements, repetitive sequence, chromatin domains, and cellular states rather than from a single dominant modifier^25^.

Together, the current data support a model in which the PWK genetic background is associated with reduced immune-associated restraint and a reorganized chromatin landscape that favors distinct malignant epithelial states under NeuT signaling. This provides a genomic explanation for how the same oncogenic driver can produce different tumor trajectories across inherited backgrounds without requiring uniformly stronger NeuT activity in PWK tumors.

## MATERIALS AND METHODS

### Study design and biological replication

Spontaneous mammary tumors were collected from female MMTV-NeuT-positive mice on the BALB/c (referred to as BALB NeuT) or (BALBxPWK) F1-derived genetic (PWK NeuT) backgrounds when tumors averaged ∼300 mm^3^. Tumor tissue was flash frozen in liquid nitrogen. Bulk RNA-seq and histone ChIP-seq analyses included five PWK NeuT and three BALB NeuT tumors, whereas single-cell multiome profiling was performed on an independent cohort of three tumors per strain. Unless otherwise indicated, positive log2 fold changes denote higher signal in PWK NeuT mice and negative values denote higher signal in BALB NeuT mice.

In the parental BALB NeuT colony, spontaneous mammary tumors develop in females at approximately 14 weeks of age, with tumor palpation beginning at 10 weeks and all mice bearing tumors by 19 weeks of age. In the PWK NeuT cohort, palpable tumors arose between 10 and 13 weeks of age and were present in all mice by 14 weeks. Tumors were monitored by palpation and caliper measurement taken two times weekly. Tumor volume was calculated as *X* × *Y*^2^/2, and tumor onset was defined as a total tumor volume of 63 mm³.

### RNA sequencing and differential expression

RNA-seq libraries were prepared using the NEBNext Ultra II RNA Library Prep Kit for Illumina (New England Biolabs) according to the manufacturer’s protocol, generating non-directional Illumina-compatible libraries.

Bulk RNA-seq reads were processed against the mm10/GRCm38 mouse genome assembly and corresponding gene annotation. Gene-level counts were filtered to remove lowly expressed features before statistical analysis. Differential expression between PWK NeuT and BALB NeuT tumors was analyzed with edgeR using biological tumors as independent samples and Benjamini-Hochberg correction to control the false-discovery rate ^26^. Principal-component analysis was performed on normalized expression values. Strain-associated genes were used for downstream pathway and regulatory analyses.

### Histone ChIP sequencing

ChIP-seq was performed for H3K4me3, H3K27ac, H3K9me3, and H4K20me3 together with matched input DNA using procedures adapted from our previously described tissue ChIP-seq workflow ^27^. Frozen mammary tumor tissue was pulverized under liquid nitrogen and crosslinked with 1% formaldehyde for 10 min at 37°C. ChIP was performed using antibodies against H3K4me3 (Millipore, 17-614), H3K27ac (Abcam, ab4729), H3K9me3 (Abcam, ab8898), and H4K20me3 (Abcam, ab9053). Reads were aligned to the mm10/GRCm38 genome assembly and filtered according to mapping-quality and duplicate-read criteria. Histone-enriched regions were identified relative to matched input, and normalized signal tracks were generated for quantitative and genome-browser analyses. Peak abundance and other strain-level summaries were calculated using tumor-level measurements.

### Expression-stratified metagene and broad-domain analyses

Genes were stratified according to bulk RNA expression. Promoters were defined as 2 kb upstream to 0.5 kb downstream of the annotated transcription start site; extended upstream intervals used for selected gene-centered analyses spanned 50 kb upstream to 5 kb downstream of the transcription start site. Gene-body metaprofiles used scaled gene bodies with flanking upstream and downstream regions. ChIP signal was normalized to matched input before tumor-level summarization. Relative redistribution was quantified from changes in the gene-body-to-promoter signal ratio.

Broad H3K9me3- and H4K20me3-associated domains were evaluated by genomic coverage and overlap between the two marks. The genome was also partitioned into fixed 1-Mb windows, and PWK-minus-BALB/c signal changes for H3K9me3 and H4K20me3 were compared using Spearman rank correlation. Tumor-level exact label permutations were used for selected comparisons where applicable.

Broad H3K9me3 and H4K20me3 domains were called with SICER using 200-bp windows, a 400-bp gap, and FDR 0.001. Fixed-window analyses used 1-Mb genomic windows and excluded windows lacking finite measurements for both marks. Strain effects were computed after averaging technical libraries within tumor. Spearman correlation quantified monotonic associations, and selected bulk comparisons used exact two-sided permutation tests over all 56 assignments of five PWK and three BALB/c tumor labels; Benjamini-Hochberg correction was used where multiple genomic features were tested.

### ChromHMM and genomic annotation

Combinatorial chromatin states were learned jointly from H3K4me3, H3K27ac, H3K9me3, and H4K20me3 using a 16-state ChromHMM model^22^. Histone-mark signal was binarized before model training, and state identities were interpreted from the learned emission probabilities rather than assigned in advance. State coverage was quantified for PWK NeuT and BALB NeuT tumors to identify strain-associated expansion or contraction of individual chromatin configurations.

Histone-enriched regions were also annotated with respect to promoters, exons, introns, intergenic regions, untranslated regions, transcription termination sites, and annotated noncoding features. Genomic-feature composition was calculated after summarizing technical measurements within biological tumors, and strain differences were expressed as changes in feature representation.

ChromHMM BinarizeBed was run in-peaks mode against the mm10 chromosome-size file, using the called regions for H3K4me3, H3K27ac, H3K9me3, and H4K20me3 as binary mark-presence inputs. A joint 16-state model was learned with LearnModel using the mm10 assembly and 200-bp bins. State labels were assigned after training from learned emission probabilities and genomic overlaps. Feature-overlap files used mm10 RefSeq gene, exon, TSS, TSS ±2 kb, TES, and CpG-island annotations; intronic and intergenic classes were derived relative to these RefSeq features.

### Repeat-centric analyses

RepeatMasker annotations corresponding to the ChIP-seq genome assembly were grouped by repeat class, family, and subfamily. Enrichment of repeat annotations within H3K9me3- and H4K20me3-associated regions was quantified from base-pair overlap relative to genomic repeat coverage. Repeat-family and repeat-territory effects were compared between PWK NeuT and BALB NeuT tumors using tumor-level chromatin measurements.

For subfamily-level analyses, repeat-associated regions were first selected on the basis of pooled H3K9me3 enrichment. H3K9me3 and H4K20me3 strain effects were then evaluated within this H3K9me3-enriched subset, avoiding direct selection on the H4K20me3 response. Thus, subfamily-level results represent H3K9me3-marked repeat regions rather than all genomic copies of each repeat subfamily. Repeat-density and joint H3K9me3-H4K20me3 change were further stratified into quintiles and related to nearby bulk RNA-seq effects.

### Single-cell multiome processing

Single-cell multiome profiling was performed on three PWK NeuT and three BALB NeuT mammary tumors using the 10X Genomics Single Cell Multiome ATAC + Gene Expression platform. Gene-expression and chromatin-accessibility libraries were generated from isolated nuclei according to the manufacturer’s protocol, following the general workflow previously described for single-cell multiome profiling of teratoma tissue^27^. Separate GEX and ATAC libraries were generated from the same nuclei.

GEX libraries were sequenced using 28 × 90-bp paired-end reads and ATAC libraries using 51 × 51-bp paired-end reads on an Illumina NovaSeq X Plus at the University of Michigan Advanced Genomics Core/Single Cell and Spatial Analysis Shared Resource. Base-call files were converted using BCL Convert v4.3.13.

Raw sequencing data were processed with Cell Ranger ARC against the mm39/GRCm39 reference, followed by multimodal analysis with Seurat and Signac using weighted-nearest-neighbor integration^19, 20^. Cells were clustered on the WNN graph and visualized using UMAP.

### Quality-control, multimodal-integration, and cell-state-annotation

Raw gene-expression and chromatin-accessibility reads were processed jointly with Cell Ranger ARC v2.1.0. Reads were aligned to the Mus musculus GRCm39 genome using the 10X Genomics refdata-cellranger-arc-GRCm39-2024-A reference package. Cell Ranger ARC called 50,369 nuclei across the six libraries before downstream filtering.

Downstream analysis was performed in R 4.2.2 using Seurat v4.3.0.1 and Signac v1.11.0. Each sample was represented as a Seurat object containing RNA and ATAC assays, and the six objects were merged. Nuclei were retained when they had 1,000-24,999 RNA counts, 5,001-69,999 ATAC counts, and less than 20% mitochondrial RNA; 32,304 nuclei passed these filters.

RNA counts were normalized and variance stabilized with SCTransform (sctransform v0.3.5), followed by principal-component analysis with 50 components. ATAC counts were transformed with Signac’s term-frequency-inverse-document-frequency procedure, all detected peaks were retained with FindTopFeatures(min.cutoff = “q0”), and singular-value decomposition was used to obtain latent semantic indexing dimensions. Weighted nearest neighbors were calculated from RNA principal components 1-50 and ATAC LSI dimensions 2-50 using FindMultiModalNeighbors with otherwise default Seurat parameters. The weighted shared-nearest-neighbor graph was clustered with the Leiden algorithm (algorithm = 3) at resolution 0.8. RNA, ATAC, and WNN UMAP embeddings were generated with RunUMAP using RNA dimensions 1-50, ATAC dimensions 2-50, or the weighted-nearest-neighbor graph, respectively; parameters not explicitly supplied to RunUMAP used the defaults in Seurat v4.3.0.1/uwot v0.1.16.

Curated cell-state annotations were generated in this study and were not assigned using an automated cell-type annotation package or external reference atlas. For each WNN cluster, we examined RNA expression profiles using cluster-level marker statistics and DotPlots generated from the study dataset. Broad lineage identities were assigned using established epithelial/luminal, basal/myoepithelial, fibroblast, endothelial, immune/myeloid, lymphoid, and adipocyte markers. Epithelial states were further characterized using Erbb-family expression and neuronal-like/axon-guidance, luminal-alveolar, secretory/inflammatory, hypoxia, interferon-response, stress-response, and cell-cycle programs. The marker genes and expression profiles supporting these curated annotations are provided in **Fig. S4**.

### Tumor-level single-cell differential analysis

Differential ATAC regions were paired with nearby or linked genes for integrative analysis. RNA-accessibility pairs were classified as concordant when gene expression and chromatin accessibility changed in the same strain direction.

RNA and ATAC count matrices were summed separately for each biological tumor within each curated cell state. A tumor-state pseudobulk sample was retained only when it contained at least 20 nuclei, and state-level testing required representation of all six tumors (three PWK and three BALB/c). Differential RNA expression and differential accessibility used edgeR v4.4.2: low-abundance features were filtered with filterByExpr using strain groups, library sizes were normalized by the trimmed mean of M values, and a design matrix of ∼ strain (BALB/c reference) was fit with the quasi-likelihood negative-binomial workflow (estimateDisp, glmQLFit, and glmQLFTest for the strainPWK coefficient). Benjamini-Hochberg FDR <0.05 and absolute log2 fold change >=0.25 defined significant features. For the **Fig. 5E** integration, differential peaks were assigned to the nearest mm39 gene and retained when the peak-gene distance was <=100 kb; pairs were called concordant when significant RNA and ATAC effects had the same sign.

### Motif activity and transcription-factor integration

Transcription-factor motif instances were identified within accessible chromatin, and chromVAR deviation scores were calculated using GC- and accessibility-matched background peaks to quantify accessibility associated with transcription-factor motifs ^21^. Motif activity was summarized by biological tumor and WNN state and compared with expression of the corresponding transcription factor.

Concordance between motif accessibility and TF RNA was treated as complementary support rather than evidence of direct transcription-factor binding.

Motifs were obtained from JASPAR2020 v0.99.10 for Mus musculus (taxon 10090; current motif versions only) and matched to mm39 peak sequences with Signac AddMotifs/motifmatchr v1.28.0 using BSgenome.Mmusculus.UCSC.mm39 v1.4.3. chromVAR v1.28.0 was run after GC-bias calculation with Signac addGCBias; GC- and accessibility-matched background peaks were generated with getBackgroundPeaks and other parameters were left at package defaults. Cell-level motif z scores were averaged within each tumor-state group. Tumor-level strain contrasts used limma v3.62.2 with a ∼ strain model and eBayes(trend = TRUE), requiring at least 20 nuclei per tumor-state and all six tumors. Motifs were significant at Benjamini-Hochberg adjusted P <0.05 and absolute PWK-minus-BALB/c mean z-score difference >=0.25. Displayed TF-motif pairs required an unambiguous motif-to-TF gene-symbol match and concordant, significant tumor-level TF RNA and motif-accessibility effects.

### Peak-to-gene associations and locus visualization

For locus-level visualization, normalized ATAC signal was displayed separately for each biological tumor together with gene models, transcription start sites, differential-accessibility regions, and peak-to-gene links on a common genomic coordinate axis. The same signal normalization and display scale were applied across strains within each locus.

Candidate peak-gene links were generated with Signac v1.11.0 using accessibility-expression covariation and mm39 gene coordinates; LinkPeaks tested peaks within 500 kb of a gene and required at least 20 cells, with package-default score and P-value calculations unless explicitly stated. Proximity-based integrations used ClosestFeature to assign the nearest mm39 gene and, for the principal RNA-ATAC pairing analysis, retained peaks within 100 kb. Links and nearest-gene assignments were combined with tumor-level edgeR RNA and ATAC effects; statistical links were retained at nominal P <0.05 and displayed only when they also met the analysis-specific concordance criteria. Coverage tracks were produced from the Signac fragment object with depth/cell-number normalization applied consistently across biological tumors, and all strain panels at a locus used a common y-axis scale.

### Trajectory inference

Malignant epithelial cells were analyzed separately using independently fitted neuronal-like and stress-secretory Slingshot trajectories. Pseudotime was interpreted as an ordering of transcriptionally and epigenetically related states rather than direct lineage ancestry.

Gene-expression dynamics along each trajectory were modeled as functions of pseudotime separately by strain. Genes displayed in **Fig. 7** were selected according to the trajectory-divergence criteria defined for the analysis, and fitted values and uncertainty were calculated along the common pseudotime axis.

Trajectory analysis used slingshot v2.14.0 and tradeSeq v1.20.0. Models included curated malignant epithelial states only. The stress-secretory model began at the Ltf-high state and included cycling, stressed, secretory/inflammatory, hypoxia/Vegfa, interferon/B2m, and Erbb2-associated states, with Spp1/Lcn2, Chil1, and hypoxia/Vegfa states evaluated as terminal branches. The neuronal-like model began at Cycling tumor epithelial A and included neuronal, Tenm3, Dpp10, axon-guidance, Erbb4, and Tshz2 states, with Dpp10, axon-guidance, and neuronal-program states evaluated as endpoints. Candidate topology was evaluated in RNA PCA dimensions 1-20, ATAC LSI dimensions 2-21, a z-scaled joint PCA-LSI space, and WNN UMAP after balancing to at most 100 cells per state and replicate. Final pseudotime used the joint RNA-ATAC latent representation, was oriented from the stated starting state, and was rescaled to 0-1. Stability was assessed by refitting after exclusion of each tumor and by 100 within-state bootstrap resamples (seed 14533). Initial gene dynamics used tradeSeq generalized additive models with six knots on a replicate-balanced subset of at most 350 cells per replicate-state and the 1,200 most variable genes. Final strain-divergence testing summarized expression in eight pseudotime-quantile bins per replicate and fit weighted mixed GAMs in mgcv with expression ∼ strain + s(pseudotime, k = 5) + s(pseudotime, by = strain, k = 5) + s(replicate, bs = ‘re’), using REML and bin cell counts as weights; reduced models omitted the strain-by-pseudotime smooth. Model-based 95% intervals were plotted. Benjamini-Hochberg correction was applied within each trajectory model. Exact strain-label inference enumerated all 20 assignments of PWK and BALB/c tumors. **Figure 7** genes were selected from the tested variable-gene set by adjusted P value, strain-curve divergence, effect magnitude, and leave-one-tumor-out stability.

### Statistics and reproducibility

Biological tumors were the unit of replication for strain-level inference. Single-cell RNA and ATAC analyses used tumor-level aggregation for inferential comparisons, whereas individual cells were used for visualization, clustering, and cell-state characterization. Unless otherwise stated, tests were two-sided and multiple comparisons were adjusted using the Benjamini-Hochberg false-discovery-rate procedure. Exact permutation tests were used for selected tumor-level analyses as specified above.

## DECLARATIONS

### Ethics approval and consent to participate

All animal procedures were approved by the Wayne State University Institutional Animal Care and Use Committee under protocol IACUC-23-06-5898 and were conducted in accordance with institutional animal-care guidelines.

### Consent for publication

All authors have read and approved the final version of this manuscript.

### Data availability

Raw and processed sequencing data have been deposited in NCBI GEO.

### Authors’ contributions

B.L.K. conceptualization, investigation, methodology, formal analysis, data curation, visualization, writing–original draft, writing–review and editing, supervision, project administration, resources, and funding acquisition; R.G. investigation and validation, including ChIP-seq experiments; J.B.J. conceptualization, investigation, supervision, project administration, resources, and funding acquisition, including mouse breeding, tumor monitoring, and mammary tumor sample collection. All authors reviewed and approved the final manuscript.

## Acknowledgements

This work utilized the Wayne State University High Performance Computing Grid for computational resources, and the Animal Model and Therapeutics Evaluation Core (AMTEC) facility.

## SUPPLEMENTAL INFORMATION

**Figure S1. Biological-replicate signal at strain-unique histone-enriched regions.**

(A-B) H3K4me3 signal centered on BALB/c-unique (A) and PWK-unique (B) enriched regions. (C,D) H3K27ac signal at BALB/c-unique (C) and PWK-unique (D) regions. (E-F) H3K9me3 signal at BALB/c-unique (E) and PWK-unique (F) regions. (G,H) H4K20me3 signal at BALB/c-unique (G) and PWK-unique (H) regions. Heatmaps show normalized signal across biological tumors over a common interval centered on the corresponding enriched regions.

**Figure S2. Repeat-subfamily and tumor-level repeat-territory effects.**

(A) Heatmap of PWK-minus-BALB/c H3K9me3 and H4K20me3 signal differences across selected H3K9me3-enriched repeat subfamilies. Positive values indicate higher signal in PWK and negative values indicate higher signal in BALB/c. (B) Tumor-level within-mark z-score heatmaps across LTR, ERV1, ERVK, satellite, and L1 territories for H3K9me3 and H4K20me3. Columns represent individual biological tumors. Source data are provided in the corresponding supplemental source table.

**Figure S3. Strain representation across the WNN atlas.**

(A) WNN UMAP displayed separately for BALB/c and PWK cells using identical embedding coordinates. (B) PWK contribution to the mean representation of each WNN cluster across the two strains. The reference line denotes equal strain representation. Cluster identities correspond to Fig. 5B.

**Figure S4. RNA marker architecture across 34 WNN clusters.**

Heatmap of cluster-associated RNA markers across the 34 WNN states. Cells are ordered by WNN cluster, strain, and biological tumor, and expression is shown as row-scaled z-score. Top annotations indicate cluster and strain. The displayed marker patterns support the curated epithelial, immune, stromal, vascular, and adipocyte state annotations used in Fig. 5.

**Figure S5. Replicate-level cis-regulatory evidence at prioritized PWK-high loci.**

(A-D) Representative PWK-high loci showing normalized ATAC coverage across BALB NeuT and three PWK NeuT tumors. Each panel includes gene annotation, transcription start site, differential-accessibility regions, genomic coordinates, and peak-to-gene associations. RNA and ATAC effect sizes and statistical support are shown within the panels. These loci provide replicate-level support for the integrated RNA-accessibility analysis in Fig. 5E.

**Figure S6. Cross-platform convergence identifies PWK-high regulatory nodes and an attenuated immune-response program.**

(A) PWK-high genes ranked by bulk RNA effect together with support from state-resolved RNA, ATAC accessibility, and histone-modification data. Point size denotes bulk RNA statistical support, color identifies the cell state contributing single-cell evidence, and the integrated score summarizes support across genomic layers. (B) Gene Ontology enrichment for genes increased or decreased in PWK tumors. PWK-low genes are enriched for interferon response, antiviral defense, antigen processing and presentation, cytokine response, and cell-killing programs, whereas PWK-high genes are enriched for cell-junction, lipid-metabolic, and differentiation-associated processes. Bar length denotes enrichment relative to the expressed-gene background, and labels indicate the number of genes contributing to each term.

