## Supplemental Figures for "PWK genetic background reprograms heterochromatin and malignant epithelial states in Neu-driven mammary tumors"

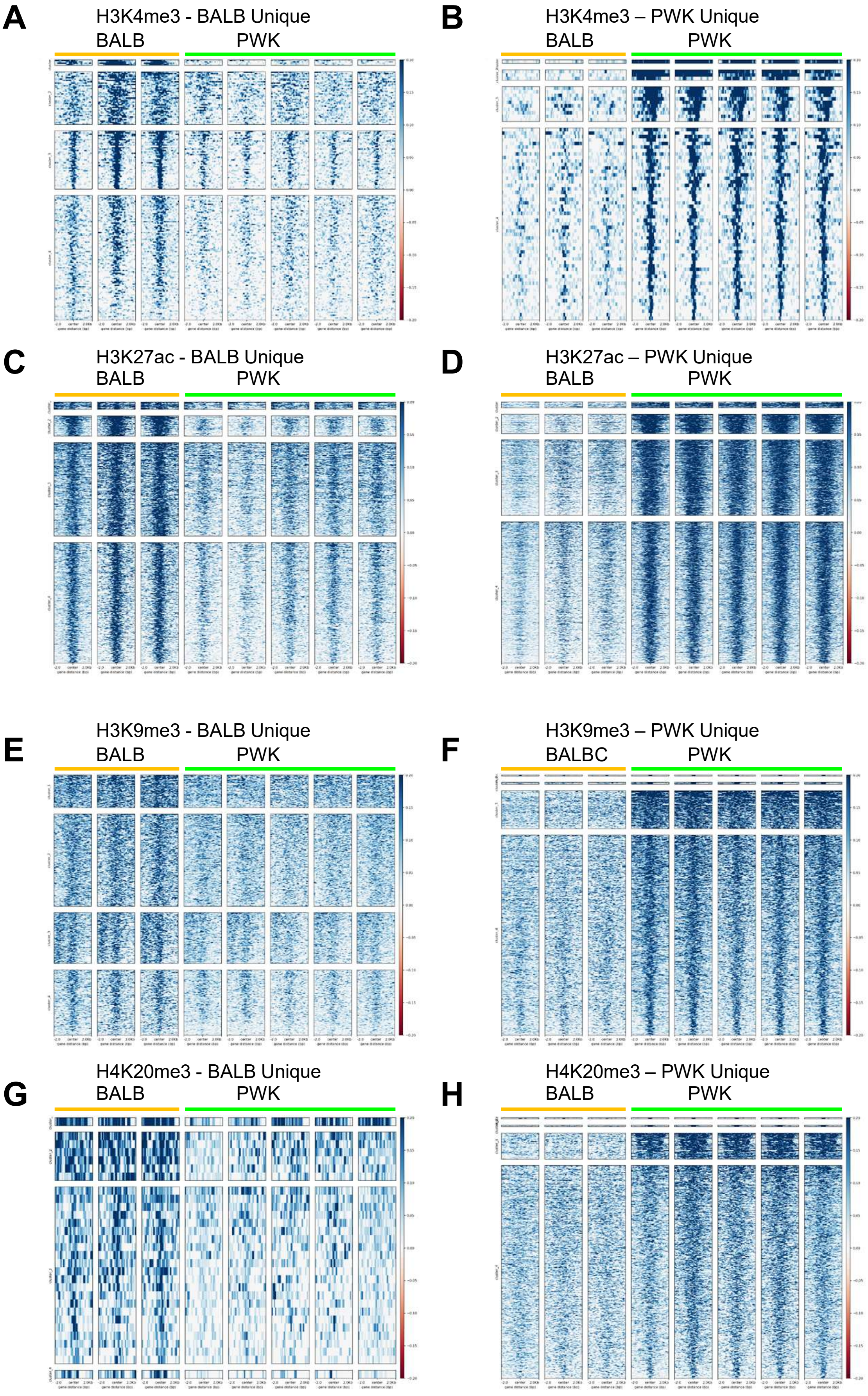
A
H3K4me3 - BALB Unique
BALB
PWK
B
H3K4me3 – PWK Unique
BALB
PWK
C
H3K27ac - BALB Unique
BALB
PWK
D
H3K27ac – PWK Unique
BALB
PWK
E
H3K9me3 - BALB Unique
BALB
PWK
F
H3K9me3 – PWK Unique
BALBC
PWK
G
H4K20me3 - BALB Unique
BALB
PWK
H
H4K20me3 – PWK Unique
BALB
PWK

**Figure S1**

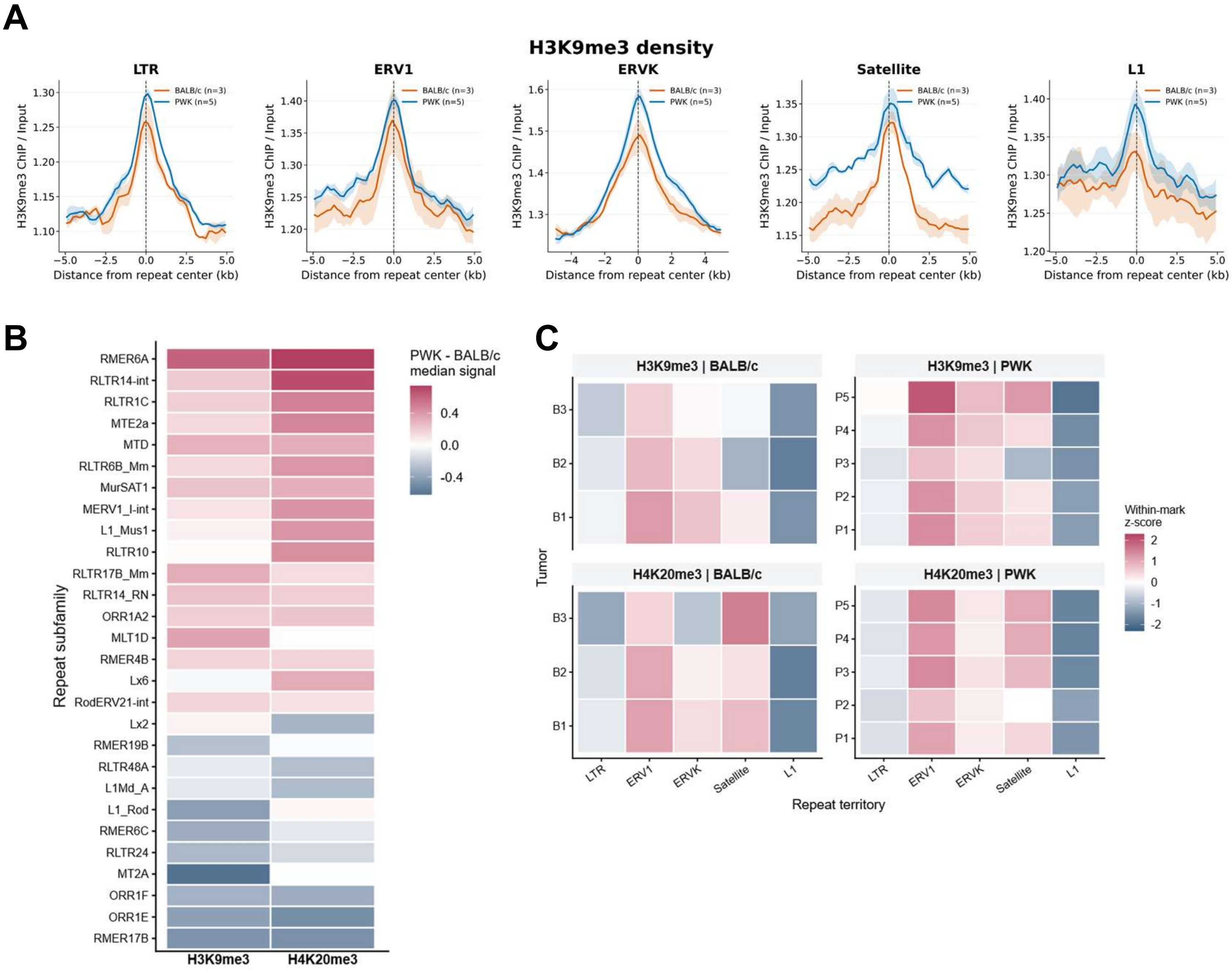
A
H3K9me3 density
LTR
ERV1
ERVK
Satellite
L1
BALB/c (n=3)
PWK (n=5)
H3K9me3 ChIP / Input
Distance from repeat center (kb)
B
PWK - BALB/c
median signal
0.4
0.0
-0.4
Repeat subfamily
RMER6A
RLTR14-int
RLTR1C
MTE2a
MTD
RLTR6B_Mm
MurSAT1
MERV1_I-int
L1_Mus1
RLTR10
RLTR17B_Mm
RLTR14_RN
ORR1A2
MLT1D
RMER4B
Lx6
RodERV21-int
Lx2
RMER19B
RLTR48A
L1Md_A
L1_Rod
RMER6C
RLTR24
MT2A
ORR1F
ORR1E
RMER17B
H3K9me3
H4K20me3
C
H3K9me3 | BALB/c
H3K9me3 | PWK
H4K20me3 | BALB/c
H4K20me3 | PWK
Tumor
B3
B2
B1
P5
P4
P3
P2
P1
LTR
ERV1
ERVK
Satellite
L1
Repeat territory
Within-mark
z-score
2
1
0
-1
-2

Figure S2

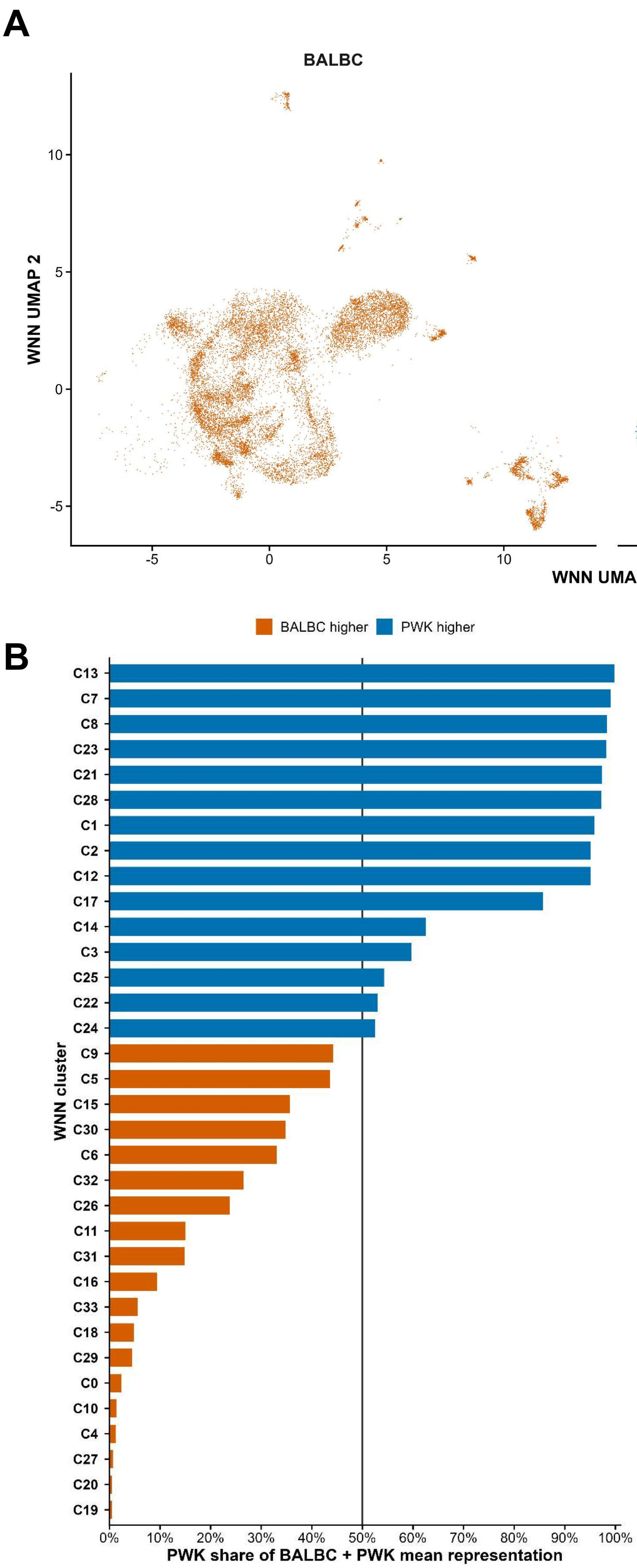
A
BALBC
PWK
WNN UMAP 2
WNN UMAP 1
10
5
0
-5
-5
0
5
10
BALBC higher
PWK higher
B
WNN cluster
C13
C7
C8
C23
C21
C28
C1
C2
C12
C17
C14
C3
C25
C22
C24
C9
C5
C15
C30
C6
C32
C26
C11
C31
C16
C33
C18
C29
C0
C10
C4
C27
C20
C19
0%
10%
20%
30%
40%
50%
60%
70%
80%
90%
100%
PWK share of BALBC + PWK mean representation

Figure S3

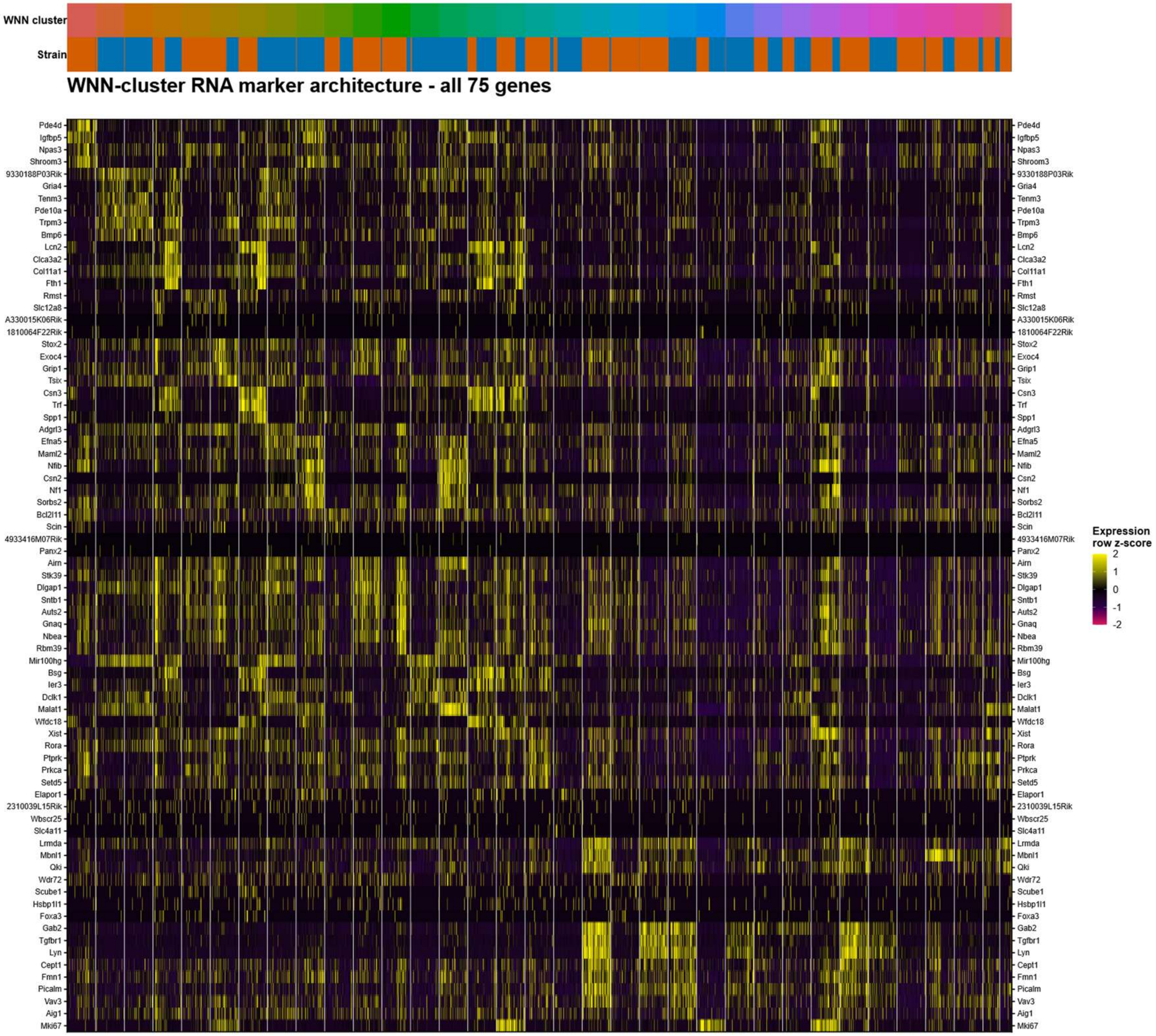
WNN cluster
Strain
WNN-cluster RNA marker architecture - all 75 genes
Pde4d
Igfbp5
Npas3
Shroom3
9330188P03Rik
Gria4
Tenm3
Pde10a
Trpm3
Bmp6
Lcn2
Clca3a2
Col11a1
Fth1
Rmst
Slc12a8
A330015K06Rik
1810064F22Rik
Stox2
Exoc4
Grip1
Tsix
Csn3
Trf
Spp1
Adgrl3
Efna5
Maml2
Nfib
Csn2
Nf1
Sorbs2
Bcl2l11
Scin
4933416M07Rik
Panx2
Airn
Stk39
Dlgap1
Sntb1
Auts2
Gnaq
Nbea
Rbm39
Mir100hg
Bsg
Ier3
Dclk1
Malat1
Wfdc18
Xist
Rora
Ptprk
Prkca
Setd5
Elapor1
2310039L15Rik
Wbscr25
Slc4a11
Lrmda
Mbnl1
Qki
Wdr72
Scube1
Hsbp1l1
Foxa3
Gab2
Tgfbr1
Lyn
Cept1
Fmn1
Picalm
Vav3
Aig1
Mki67
Expression row z-score
2
1
0
-1
-2

Figure S4

**A**

### Sema7a locus (PWK-high)

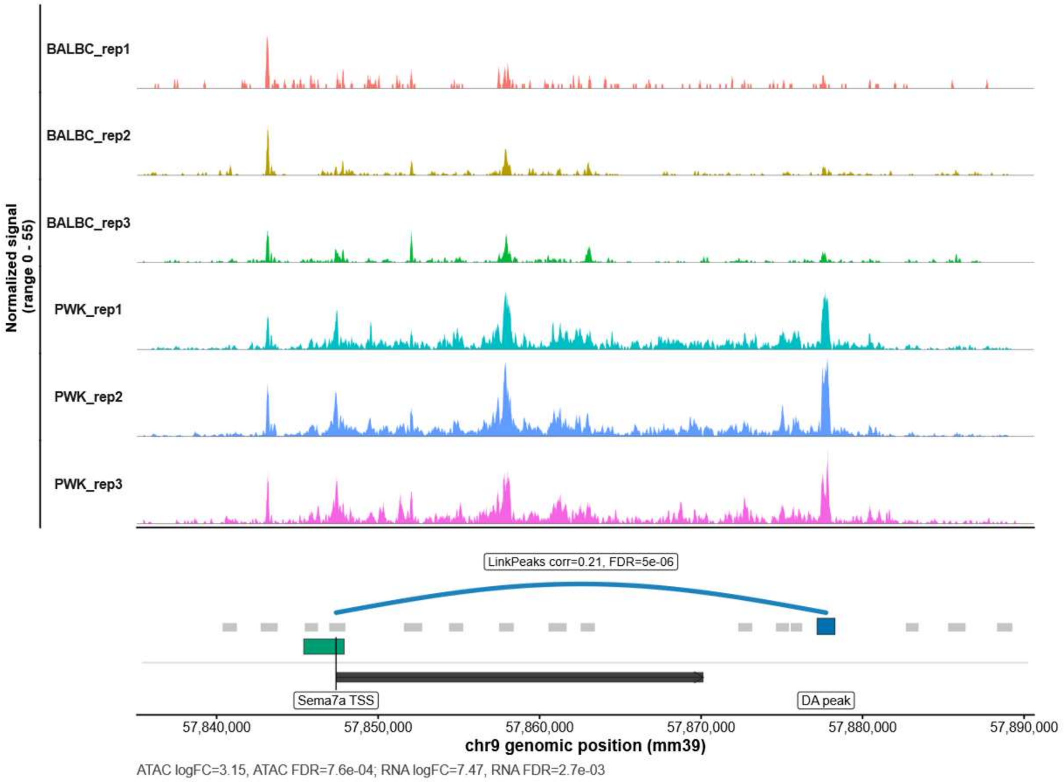


**B**

### Frem1 locus (PWK-high)

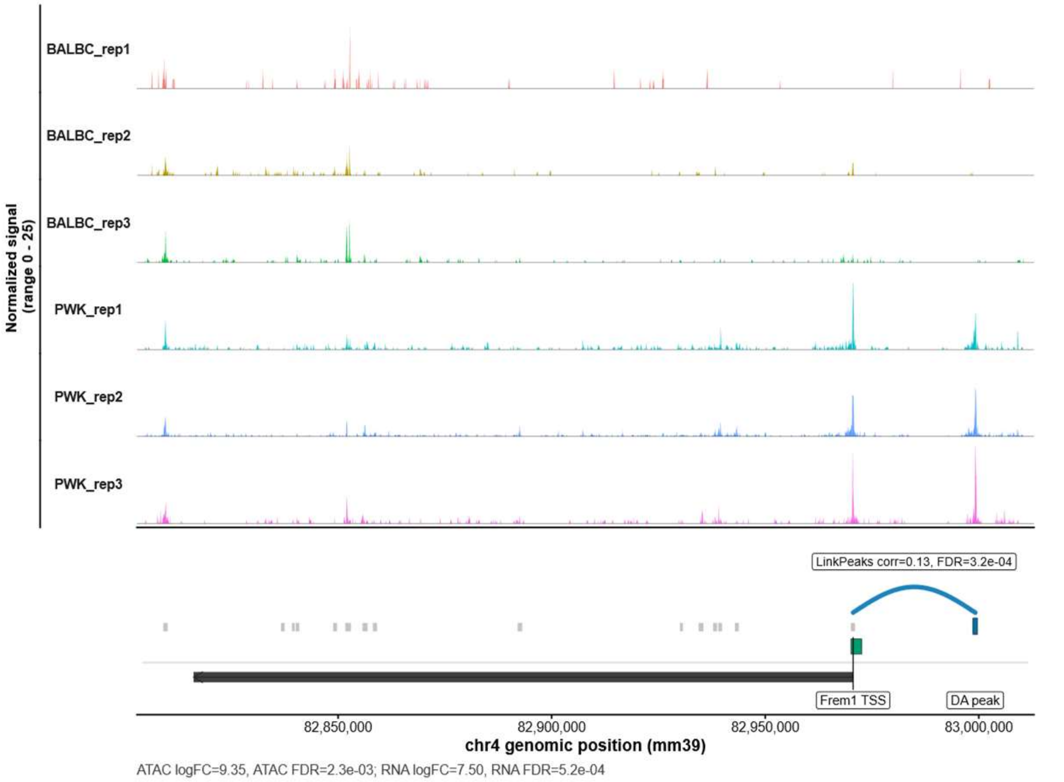


**Figure S5**

**C** Tcerg1l locus (PWK-high)

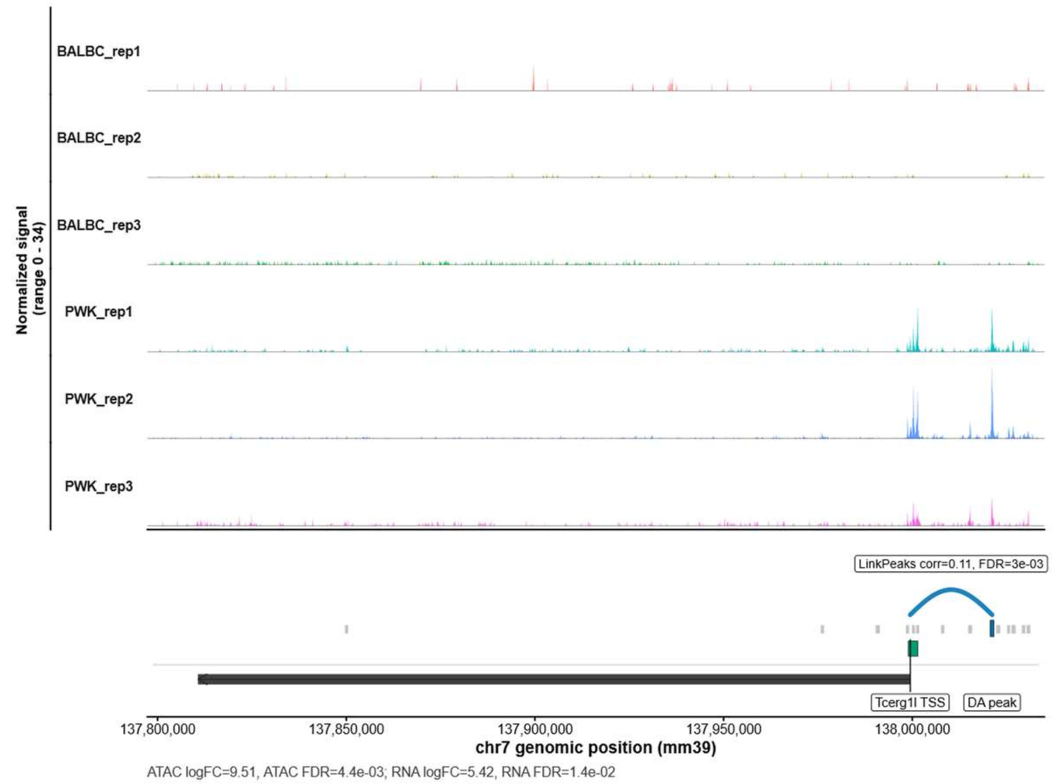


**D** Elapor2 locus (PWK-high)

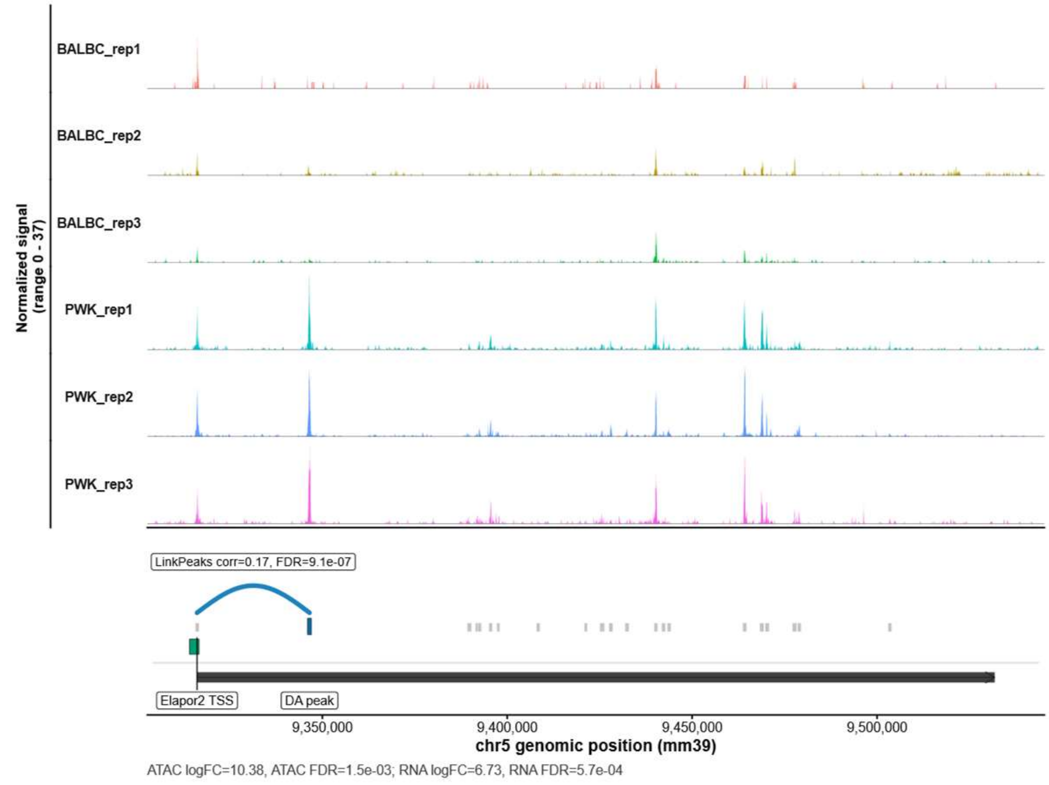


**Figure S5**

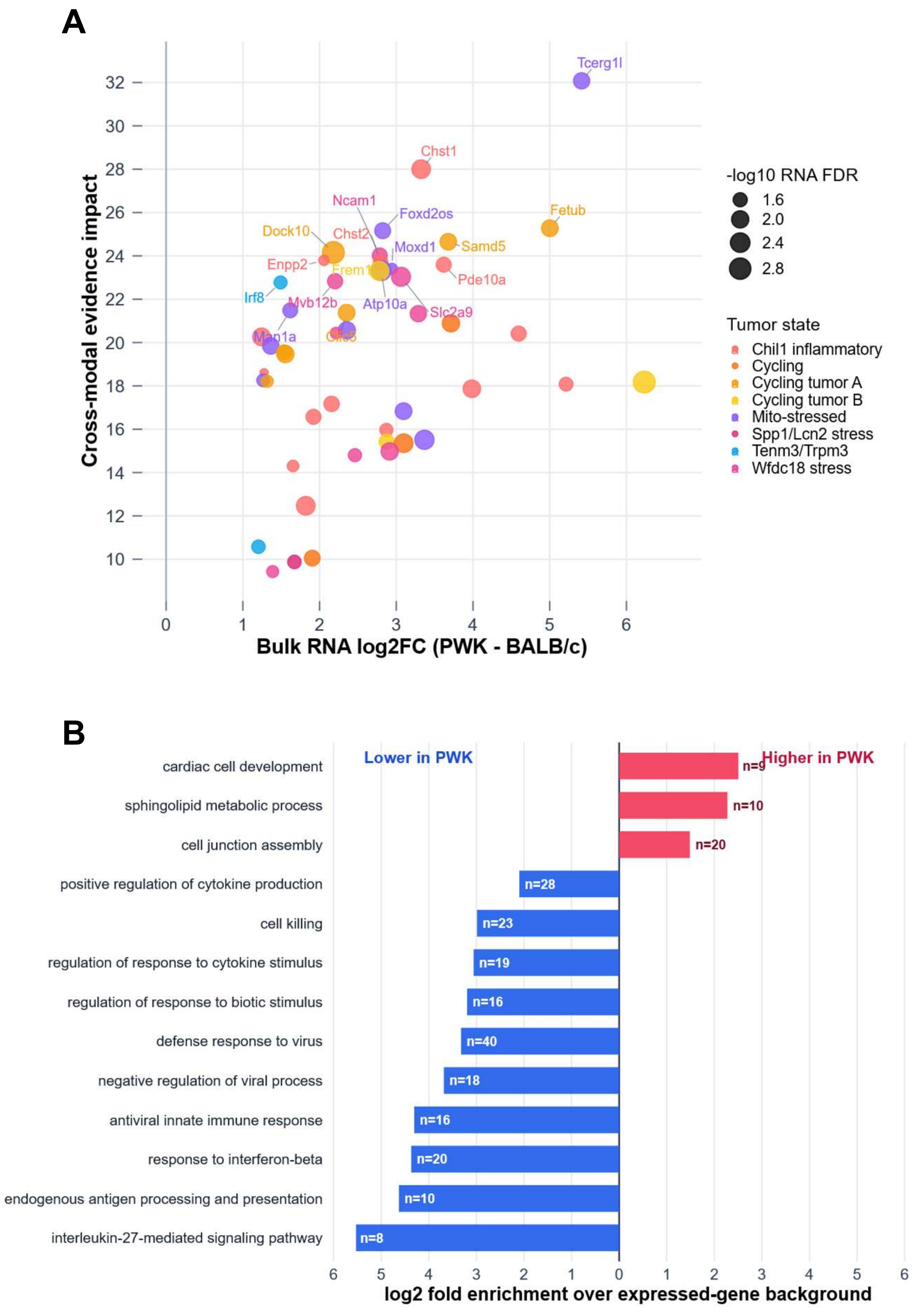

A
Cross-modal evidence impact
Bulk RNA log2FC (PWK - BALB/c)
Tcerg1l
Chst1
Ncam1
Foxd2os
Fetub
Dock10
Chst2
Moxd1
Samd5
Enpp2
Frem1
Pde10a
Irf8
Mvb12b
Atp10a
Slc2a9
Man1a
-log10 RNA FDR
1.6
2.0
2.4
2.8
Tumor state
Chil1 inflammatory
Cycling
Cycling tumor A
Cycling tumor B
Mito-stressed
Spp1/Lcn2 stress
Tenm3/Trpm3
Wfdc18 stress
B
Lower in PWK
Higher in PWK
cardiac cell development
sphingolipid metabolic process
cell junction assembly
positive regulation of cytokine production
cell killing
regulation of response to cytokine stimulus
regulation of response to biotic stimulus
defense response to virus
negative regulation of viral process
antiviral innate immune response
response to interferon-beta
endogenous antigen processing and presentation
interleukin-27-mediated signaling pathway
n=9
n=10
n=20
n=28
n=23
n=19
n=16
n=40
n=18
n=16
n=20
n=10
n=8
log2 fold enrichment over expressed-gene background


**Figure S6**
